# A high-throughput search for intracellular factors that affect RNA folding identifies *E. coli* proteins PepA and YagL as RNA chaperones that promote RNA remodeling

**DOI:** 10.1101/2024.06.04.597434

**Authors:** Alejandra Matsuri Rojano-Nisimura, Lucas G. Miller, Aparna Anantharaman, Aaron T. Middleton, Elroi Kibret, Sung H. Jung, Rick Russell, Lydia M. Contreras

**Affiliations:** Department of Molecular Biosciences, The University of Texas at Austin, Austin, Texas 78712, United States; McKetta Department of Chemical Engineering, The University of Texas at Austin, Austin, TX 78712, United States; Department of Biomedical Engineering, The University of Texas at Austin, Austin, Texas 78712, United States

## Abstract

General RNA chaperones are RNA-binding proteins (RBPs) that interact transiently and non-specifically with RNA substrates and assist in their folding into native state. In bacteria, these chaperones impact both coding and non-coding RNAs and are particularly important for large, structured RNAs which are prone to becoming kinetically trapped in misfolded states. Currently, due to the limited number of well-characterized examples and the lack of a consensus structural or sequence motif, it is difficult to identify general RNA chaperones in bacteria. Here, we adapted a previously published *in vivo* RNA regional accessibility probing assay to screen genome wide for intracellular factors in *E. coli* affecting RNA folding, among which we aimed to uncover novel RNA chaperones. Through this method, we identified and validated eight proteins whose deletion gives changes in regional accessibility within the exogenously expressed *Tetrahymena* group I intron ribozyme. Further, we purified and measured *in vitro* properties of two of these proteins, YagL and PepA, which were especially attractive as general chaperone candidates. We showed that both proteins bind RNA and that YagL accelerates native refolding of the ribozyme from a long-lived misfolded state. Further dissection of YagL showed that a putative helix-turn-helix (HTH) domain is responsible for most of its RNA binding activity but only the full protein shows chaperone activity. Altogether, this work expands the current repertoire of known general RNA chaperones in bacteria.

## Introduction

The biological functions of many RNAs require them to fold into specific conformations. RNA folding is a complex, hierarchical process in which initial chain compaction and local structure formation often result in non-native structures or contacts that become fixed by tertiary contacts to generate misfolded conformations (**1, 2**). These misfolded states can be long-lived, with large free energy barriers for refolding to their corresponding native states (**3**). General RNA chaperones are RNA-binding proteins (RBPs) that bind RNA transiently and with low specificity (affinities typically in the µM range) to resolve misfolded conformers by promoting local unfolding. Upon protein-RNA complex dissociation, the RNA has an additional opportunity to fold to its native, functional state (**4, 5**).

Known examples of general RNA chaperones in *E. coli* include the cold-shock domain protein, CspA, and the StpA protein. CspA is a relatively small (7.4 kDa) protein and a member of the OB-fold superfamily (**6**). OB-fold proteins share a characteristic five-stranded β-barrel structure with two positively charged regions surrounding an exposed aromatic patch, allowing intermediate (µM range) nucleic-acid binding (**7**). Specifically, CspA binds single-stranded RNA (ssRNA) and acts as an RNA chaperone during cold-shock (when intracellular CspA concentration is about 10^−4^ M) by resolving stem loop structures within nascent mRNAs and allowing for transcript elongation (**8, 9**). In the case of StpA, this protein has been shown to have annealing and strand displacement activity *in vitro* and to be capable of increasing the frequency and/or lifetime of local unfolding events within structured RNAs (**10**). StpA is also a small (15.3 kDa) protein and is composed of two structural domains. Intermediate nucleic acid binding (µM range affinity) occurs via electrostatic interactions between the RNA and the positively charged C-terminal domain (CTD) of the protein (**11**). Other proteins with RNA chaperone activity in *E. coli* include ribosomal proteins, with S12 being the best characterized example (**12, 13**), and ATP-dependent general RNA remodelers from the DEAD-box helicase family, like SrmB, CsdA and RhlE (**reviewed in 14**).

General RNA chaperones are both structurally and functionally diverse. Nevertheless, an emerging theme is that, in addition to their RNA chaperone functions, several of these proteins have specialized functions that involve binding to DNA substrates. For instance, CspA acts as a transcriptional enhancer by recognizing a single-stranded CCAAT motif and promoting the transcription of genes such as *hns* and *gyrA* (**15**). Likewise, StpA, a paralog of the HN-S global regulator that participates in DNA condensation, binds curved DNA and has been proposed to serve as a molecular *hns* backup (**16, 17**). These findings suggest that additional DNA-binding proteins (DBPs) might also moonlight as RNA chaperones. However, identification of other DBPs (or proteins in general) capable of remodeling RNA substrates is limited by the lack of shared structural or sequence motifs; thereby making it infeasible to predict RNA chaperone functions using bioinformatics.

To identify cellular factors that promote RNA folding, we have adapted the *in vivo* RNA Structural Sensing System (iRS^3^) assay previously developed by our group (**18, 19**). The iRS^3^ uses sequence-specific, user-designed antisense RNA (asRNA) probes against an RNA of interest by coupling hybridization of the probe to activation of a downstream GFP fluorescence reporter sequence. As such, this assay allows the evaluation of changes in local RNA structures and the identification of potential functional regions (such as those that participate in RNA-protein interactions) (**20**). We previously used this iRS^3^ assay to profile folding of the *Tetrahymena* group I intron ribozyme using a collection of sequence-specific asRNA probes and benchmarked our results to those obtained by *in vivo* DMS probing (**18**).

Here, we coupled the iRS^3^ assay to a Tn5 transposon library and identified genes that, when knocked out, give changes in the accessibility of a key region of the *Tetrahymena* group I ribozyme when it is expressed in *E. coli*. This ribozyme was selected due to its extensive structural characterization *in vitro,* which includes cryo-EM resolved structures of its native and non-native states and chemical probing structural maps (**22–27**), and its known retained activity when expressed in *E. coli*. Through this approach, we identified two genes, *yagL* and *pepA*, that may encode RNA chaperones. Supporting this hypothesis, we found that purified PepA protein, shown previously to interact functionally with DNA (**28**), can also bind single-stranded RNA (ssRNA) and double-stranded RNA (dsRNA). Further, the purified YagL protein, which was previously uncharacterized, accelerates native refolding of a long-lived misfolded conformation of the *Tetrahymena* ribozyme and binds both ssRNA and dsRNA. Via homology modelling and deep learning, we identified a predicted helix-turn-helix (HTH) RNA-binding domain in the C-terminus region of YagL and showed, using protein truncations, that this predicted HTH region is primarily responsible for the RNA-binding activity of YagL. In contrast, both the N- and C-terminal domains are required for its chaperone role *in vitro*, suggesting additional functional roles for the N-terminus of YagL. Together, our results expand the repertoire of known RNA chaperone proteins in *E. coli* and introduce a methodology to investigate their influence in RNA folding *in vivo*, which could be applicable to other RNA chaperones and other RNA molecules.

## Materials and Methods

### Plasmids and strains

A detailed list of all strains, plasmids and primers used in this study can be found under **Supplementary Table S1.**

### EZ-Tn5 transposition and library preparation

A DNA fragment containing tetR was PCR amplified from pACYC184 (**29**). pCML375 was constructed by inserting this fragment into the EZ-Tn5-carrying plasmid pMOD<MCS> (Lucigen). The transposon DNA was purified by digesting pCML375 using PvuII-HF (NEB), and transposomes were generated following the manufacturer’s instructions. 1 U/μL of the EZ-Tn5 transposome was used for electroporation into *E. coli* MG1655 as previously described in (**30**).

After transformation, recovered cells were grown on eight large plates (150x15 mm.) containing Luria-Bertani (LB) medium (Benton-Dickenson and Company) supplemented with 10 μg/mL tetracycline (Amresco). To ensure good library coverage, each plate was partitioned into 68 sections and 20-30 colonies were collected from each partition.

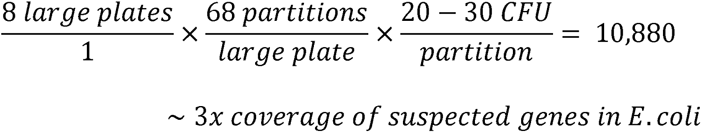

A previously published toehold-mediated reporter plasmid containing a tetracycline-inducible antisense RNA (asRNA) targeting the P3 and P4 domains of the *Tetrahymena* ribozyme (herein Probe 3) fused to GFP (**18, 31**), as well as the ribozyme target transcript under a pBAD promoter (sequence included in **Supplementary Table S2**) was transformed into the isolated colonies by chemical transformation (this plasmid is referred to as pCML2533 in **Supplementary Table S1**). In short, cells from 20-30 colonies were treated with CaCl_2_ to make them competent and were then transformed with the pCML2533 plasmid and plated in LB-agar supplemented with 10 μg/mL tetracycline and 50 μg/mL kanamycin (Amresco). From the resulting plates, library collections were assembled by collecting transformed colonies directly from the plates by resuspending them in ∼ 500 μL of fresh liquid LB media. Five different library collections were obtained, each containing recovered cells from ∼14 plates.

Cells were subcultured to prepare 25% (v/v) glycerol stocks which were stored at -80°C.

### Screening of the transposon library and FACS sorting

A 500 µL glycerol cryo-stock aliquot was thawed completely and used to inoculate 20 mL of LB medium supplemented with 10 μg/mL tetracycline and 50 μg/mL kanamycin (Amresco). Cultures were incubated at 37°C and 120 rpm until they reached target OD_600 ∼ 0.15_. At that point, the *in vivo* reporter was induced by adding 800 μL of 20% arabinose (final concentration 0.8%) and 20 μL of 100 mg/mL anhydrotetracycline (aTc) (final concentration 100 ng/μl). Prior to sorting, cultures were incubated for 2.5 hours.

Cell sorting was performed using a FACSAria IIIu cell sorter (Becton Dickinson) equipped with a blue solid-state laser (488 nm excitation) using a 100 μm nozzle at a sorting rate of ∼300 events/s and a sorting efficacy between 75 and 90%. Settings were adjusted and the data was visualized using the FACSDiva software (BD and Company). Fluorescence scatter was used to sort cells into four different populations based on their GFP fluorescence intensity (GFP-A). The area of interest was determined for high fluorescence expressing cells by comparing the scatter distribution of the transposon library relative to wild type *E. coli* MG1655 cells.

Isolates from the region of interest corresponded to 1.1-1.7% of the total population. Cells were sorted on agar plates containing LB with 10 μg/mL tetracycline and 50 μg/mL kanamycin (Amresco) and 377 different CFU were obtained. For storage, glycerol cryo-stocks were made for each isolate and were stored at -80°C.

### Validation of isolated mutants after FACS by 96-well plate reader screens

To validate the outcome of the high-throughput screen, colonies were screened again using a Cytation3 plate reader in a 96-well plate. Colonies corresponding to the 377 isolates obtained after Fluorescence Activated Cell Sorting (FACS) were re-grown in 5 mL overnight cultures supplemented with 10 μg/mL tetracycline and 50 μg/mL kanamycin (Amresco). The next morning, 96-well polystyrene plates (Greiner) were filled, with 300 μL of fresh LB broth and 0.5% kanamycin (Amresco) and inoculated with 3 μL of overnight culture (6 wells per overnight, corresponding to three technical replicates for induced and uninduced conditions). After two hours, 0.15 μL of 100 mg/mL aTc (final concentration 100 ng/μl) were added to the wells. 6 μL of 40% arabinose (final concentration 0.8%) were added to half of the wells, corresponding to the induced condition. After 5 hours of incubation, final OD_600_ measurement and fluorescence readings were collected. Sixty-four isolates were selected, using a fluorescence ratio log2 fold-change cutoff of >0.5 and a p-value of <0.05, for follow-up identification and characterization experiments. A schematic overview of the screening and validation process after FACS is shown in **Supplementary Fig. S1**.

### Whole-genome sequencing and identification of transposon insertions

Overnight cultures were started for each of the 64 library isolates using 5 mL of LB medium supplemented with standard amounts of tetracycline. These cultures were pooled together into eight different subcultures. To this purpose, 500 μL of overnight culture (62.5 µL per isolate) were used to seed 500 mL of fresh LB medium (1:1000 dilution). Cultures were grown to saturation and used to make glycerol cryo-stocks to be stored at -80°C.

For each subculture, genomic DNA (gDNA) was extracted using a QIAamp UCP DNA Micro Kit (QIAGEN) and following manufacturer’s instructions. Total yields were ∼6-8 µg of genomic DNA per pool. gDNA was extracted from wild type *E. coli* MG1655 cells as a control. Genomic DNA was submitted to the GSAF core facility (UT Austin) for library prep and sequencing. The samples were analyzed for quality check using a bioanalyzer (Agilent). DNA libraries were prepared using standard Illumina kits and were run using a MiSeq sequencer (Illumina) in a 2x250 paired-end scheme.

The following computational pipeline was used to identify transposon insertions: (i) quality control checks were performed using fastqc (**32**). (ii) adapter sequences were trimmed using CUTADAPT (**33**). (iii) the trimmed sequences in FASTQ format were used as inputs for the Transposon Insertion Finder (TIF) program (**34**). The 19-bp mosaic end (ME) sequence (*5’-CTG TCT CTT ATA CAC ATC T - 3*’) recognized by the Tn5 transposase and its reverse complement were used as head and tail end sequence inputs for the TIF program. Additionally, the length of the resulting transposon site duplication (TSD) was set to 9 bp. The resulting fasta file, containing all identified sequences flanking a transposon sorted by TSD, was used to map the transposon insertion start and end positions by BLAST search against the *E. coli* K-12 MG1655 reference genome.

In total, 31 different transposon insertions were mapped using this approach **(Table 1).**

**Table 1.**
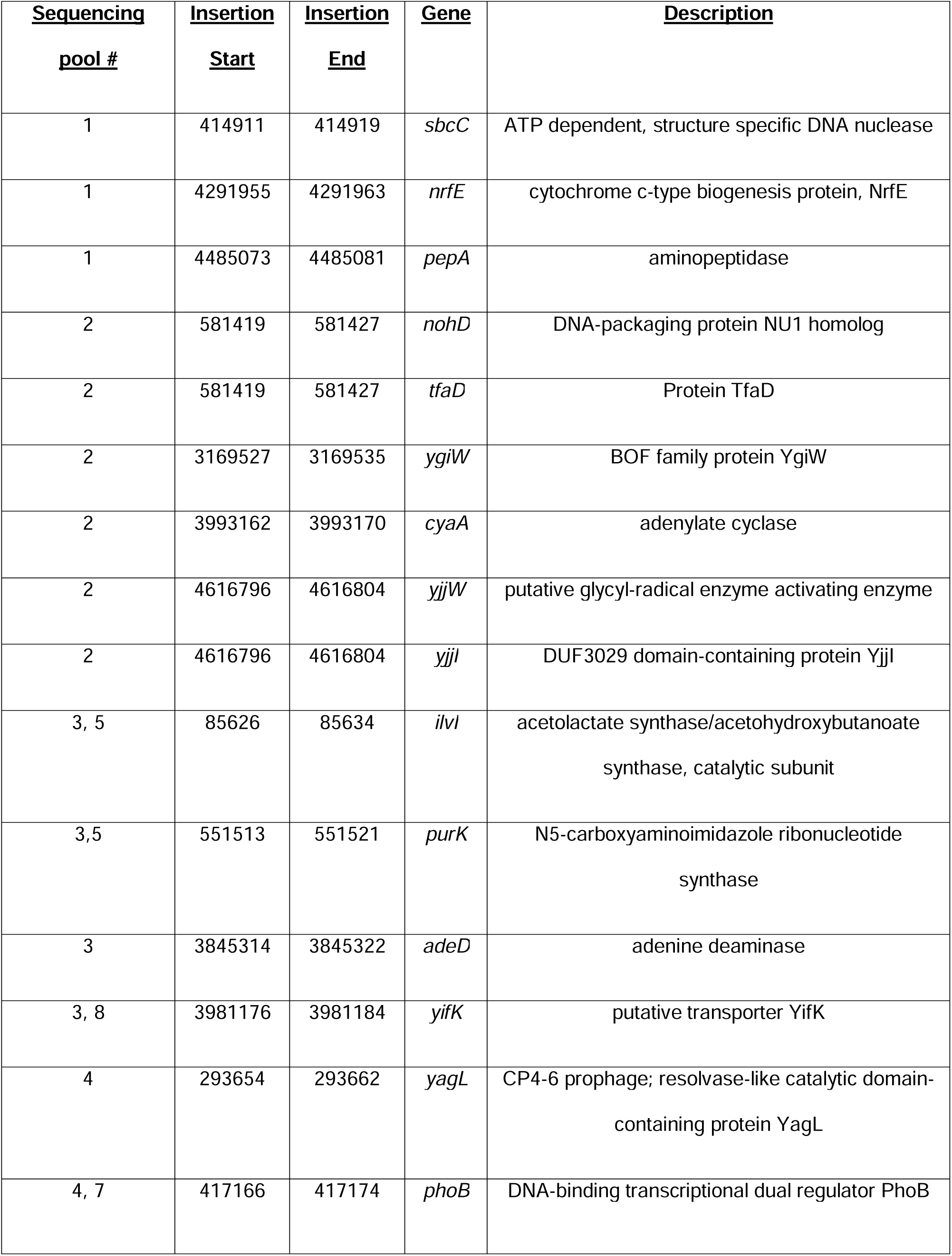
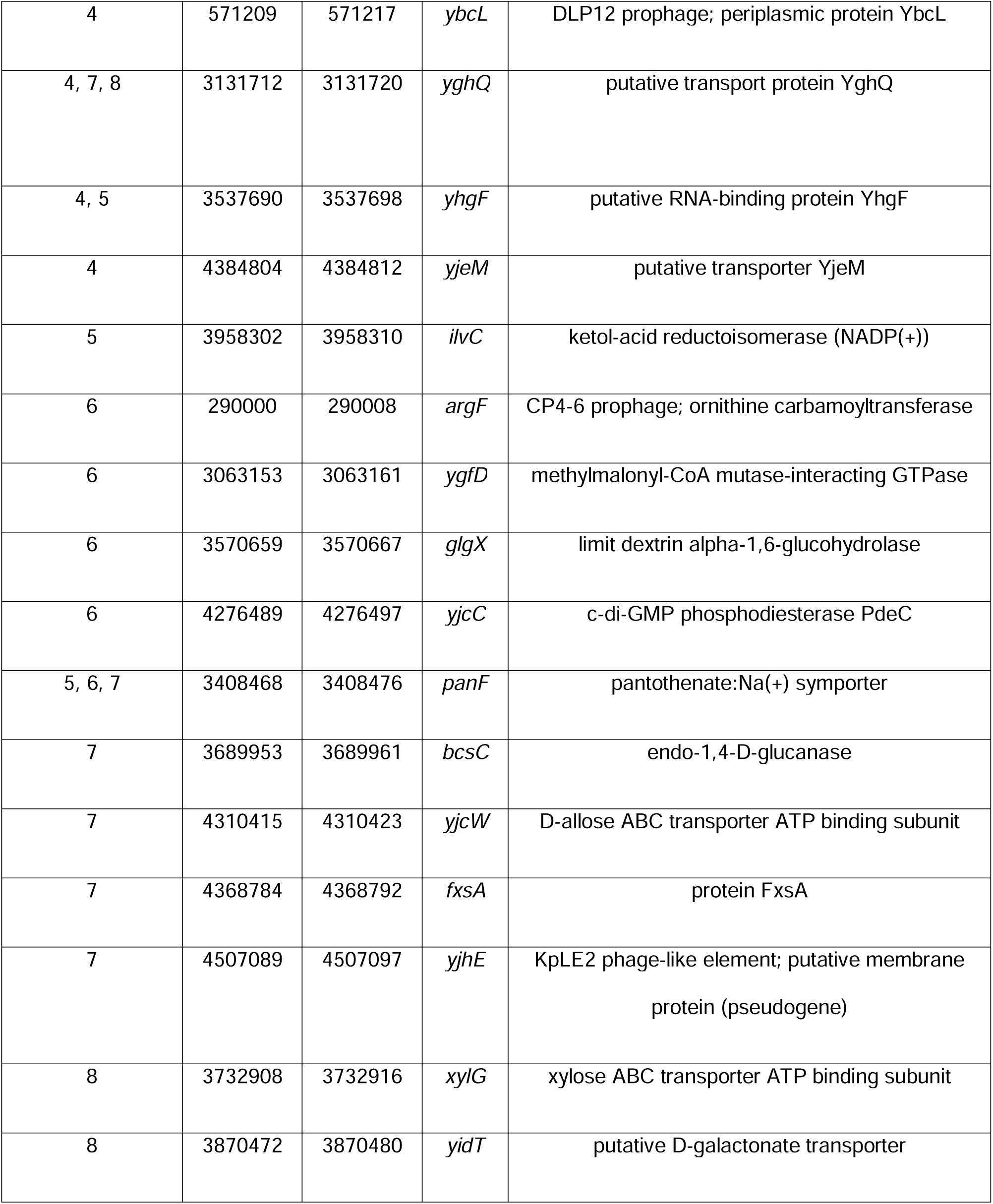
List of genes identified through whole-genome sequencing (WGS). 31 different transposon insertions were identified after cell sorting and sequencing. The sequencing pool number corresponds to the number of sample(s) in which the insertion was mapped out of the eight different pools of isolates for which gDNA was extracted. Insertion start and insertion end are the bp positions from the *E. coli* reference genome at which the transposon start and end sequence was mapped. Gene descriptions were obtained from Genbank.

### Fluorescence measurements and regional accessibility mapping using the in vivo RNA structural sensing system (iRS^3^)

Fluorescence measurements were performed by adapting a previously published protocol (**19**). In short, *E. coli* cells (*i.e.,* genomic mutants of potential chaperone genes identified by transposon screening, **Table 1**), were transformed with the toehold-mediated reporter plasmid carrying Probe 3 (see **EZ-Tn5 transposition and library preparation** section above). Additionally, strains were transformed in parallel with two previously published control probes (**19**): “p-O-iRS3GG-scramble”, which encodes for a random 15 nt asRNA probe that does not have complementarity with the genome of *E. coli* and should thus remain in an RBS-sequestered conformation and generate only background, low-end GFP signal, and “p-O-iRS3GG-open”, which is based on the scramble probe but the cis-blocking region that sequesters the RBS is mutated such that the probe is always in a free, open conformation and should generate the max, high-end of GFP signal. These control probes were used in every experiment to validate the induction of the probes upon chemical addition and to assess the fluorescence range of the experiment.No major variations were observed in the fluorescence of these control probes between experimental runs (**Supplementary Fig. S4**). When fluoresnce of the controls was not replicated, the results from those runs were not considered.

Transformed strains harboring the reporter plasmids were grown overnight at 37°C and 120 rpm. 50 μL of overnight culture were used to inoculate 5 mL of fresh LB medium (1:100 dilution) supplemented with 50 μg/mL of kanamycin (Amresco). Two tubes with fresh medium were seeded per overnight culture. Cultures were allowed to grow until they reached target OD_600_ ∼ 0.15. At that point, 5 μL of anhydrotetracycline (aTc) (final concentration 100 ng/μL) were added to the cultures to induce expression of the asRNA probe. This condition represents the target-uninduced control (herein after referred to as uninduced for simplicity). Additionally, 200 μL of 20% arabinose (final concentration 0.8%) were added to one of the culture pairs for induction of the target RNA (ribozyme) (induced condition).

GFP fluorescence was measured 2.5 hours post-induction on a FACSCalibur flow cytometer (Becton Dickinson) equipped with a 488 nm argon laser and a 530 nm FL1 logarithmic amplifier. Sample data were collected using the CellQuest Pro software (BD and Company) with a user-defined gate. Fluorescence measurements were collected from ∼250 000 cells per sample and analyzed using Microsoft Excel (Microsoft for Windows). The medians of uninduced (asRNA probe only) and induced (asRNA probe + ribozyme) samples were used to calculate the fluorescence relative ratio of each biological replicate tested.

### Protein purification

*stpA* was cloned into the *BamHI* and *SacI* sites of pET-21a (+) via Gibson assembly using pCYFP (**37**) as a starting backbone and through the addition of homology arms to the stpA sequence by PCR. The resulting plasmid (pCML2868) was used to purify StpA-H6 (containing six additional His residues at the C terminus) via Ni-NTA pull downs as described in (**38**). The same cloning approach was used to generate plasmids for the purification of PepA-H6 (pCML2870) and YagL-H6 (pCML3388). Successful protein expression and band sizes were confirmed by Coomassie blue staining **(Supplementary Fig. S2)**. After purification, protein concentrations were determined via Bradford assay using bovine serum albumin (BSA) as a standard. We confirmed that PepA was functionally active after purification by performing a peptidase activity assay (see **Supplementary Methods**). The activity for our purified PepA was determined to be ∼73.1 nKat/mg of protein **(Supplementary Fig. S3).** Previous determinations using this method have found the activity of PepA to be 30-150 nKat/mg of protein (**39**).

Gibson assembly was also used for the generation of YagL domain truncations (**40**). Briefly, primers were designed to amplify the YagL coding region from pCML3388 corresponding to the HTH domain (187–232) or the full-length protein without this region (1–186) with homology arms for insertion between StyI and XhoI sites of pET21a (+). These inserts were ligated into the pET21a (+) backbone to form YagL_HTH (pCML3739) and YagL_dHTH (pCML3725). Protein expression, purification, and quality checks were performed as described above. Protein purity was further verified by LC-MS/MS. Samples were digested with trypsin, desalted, then run on the Dionex LC and Orbitrap Fusion 2 for LC-MS/MS with a 2-hour run time. Raw data files were analyzed using Proteome Discoverer version 2.5 and Scaffold version 5. The intended his-tagged purified proteins were the most abundant proteins detected via mass spectrometry. No proteins were detected within 10-fold of the Normalized Quantitative Value for each purified protein sample.

### Northern blotting analysis

To measure steady state levels of the iRS^3^ transcript, RNA was extracted from cells expressing the iRS^3^ reporter 3h post-induction using the Direct-zol™ RNA MiniPrep kit (Zymo). After extraction, the RNA was subjected to Northern Blot analysis as described in (**35, 36**). The iRS^3^ transcript was blotted using probe I (*5’-GCCCATTAACATCACC - 3*’). For the *in vivo* assays, the iRS^3^ transcript is expressed from a pLtetO promoter.

### In vitro RNA-Protein binding assays

Filter binding assays were performed by adapting previously published protocols (**10, 41, 42**). Two 5’-end dephosphorylated oligonucleotides corresponding to a 21-mer sequence (*5*′*-AUGUGGAAAAUCUCUAGCAGU-3*′, herein “21R+”), and the complementary sequence (*5*′*-CUGCUAGAGAUUUUCCACAU-3’,* herein “21R-“), previously published in (**10**), were custom synthesized by Genelink. An additional short RNA hairpin (*5’-GCTCTAGAGCATTATGTTCAGATAAGG-3’*, herein “hairpin”), previously published in (**10**) was custom synthesized by IDT. The P^32^ end-labeled oligonucleotides were prepared as described in (**43**). Binding reactions were carried out by incubating 50 fmol of labeled RNA oligonucleotide with increasing concentrations of the respective protein (0 – 10 µM) for 10 min at room temperature in binding buffer (75 mM Tris-HCl pH 7.5, 0.4 mM spermidine, 0.1LmM MgCl_2_, 50LmMLNaCl, 0.5LmMLEDTA, 0.25LmMLDTT and 6% glycerol; 60 µL total reaction volume).

After incubation, 50 μL of the binding mixture reactions were applied onto a Bio-Dot apparatus (BioRad) assembled with membranes pre-equilibrated with binding buffer without salts (top: nitrocellulose membrane (0.45 μm pore size; BioRad); bottom: Nylon N+ membrane (0.2 μm pore size; Amersham/Cytiva) and washed twice with 100 μL binding buffer. The membranes were placed on Whatman chromatography paper (Amersham/Cytiva) to air dry for ∼10 minutes and then covered with saran wrap. The membranes were exposed to a phosphor screen (GE Healthcare) for 1 hour. Following exposure, the phosphor screen was imaged using a Typhoon Phosphorimager.

### Kinetic chaperone activity assays

The activity of candidate chaperone proteins was evaluated using a previously published *in vitro* two-step catalytic activity assay (**44, 45**). In this assay, catalytic activity is used as a readout to monitor native state folding of the *Tetrahymena* ribozyme.

Materials were prepared as described in (**46**) and measurements were performed with slight modifications. Specifically, the ribozyme (0.15 µM) was incubated in 20 mM Na-MOPS (pH 7.0) and 5 mM MgCl_2_ for 6 min at 25°C to give a population of predominantly misfolded ribozyme and then placed on ice. Folding reactions were initiated by the addition of 6 µL of purified enzyme (*i.e.,* StpA, YagL, PepA, etc.) at different concentrations (or the equivalent volume of chaperone storage buffer [20 mM Tris-Cl (pH 7.5), 500 mM KCl, 1 mM EDTA, 0.2 mM DTT, and 50% glycerol (vol/vol); 6 µL]). Refolding was monitored at 25°C. At various time points, reaction aliquots (2 µL) were transferred to a folding quench solution [2 µL of 95 mM MgCl_2_, 1 mM guanosine, 1 mg/ mL proteinase K and 20 mM Na-MOPS (pH 7.0)] to stop the folding reaction and create the necessary conditions for the subsequent substrate cleavage reaction. Quenched aliquots were kept on ice until the cleavage reactions were initiated.

During the catalytic step, substrate cleavage reactions were initiated by adding 1 µL of trace 5’-^32^P-labeled substrate oligonucleotide (∼ 20,000 dpm/ µL) to the folding reaction aliquots. This substrate (*5’-CCCUCUA_5_-3’*, abbreviated rSA_5_) mimics the 5’-splice-site junction and is cleaved by the ribozyme to give a shorter radiolabeled product (*5’-CCCUCU -3’*) (**47**). Cleavage reactions were stopped after 2 min with 2-fold excess of an EDTA-containing gel-loading solution [72% formamide (vol/vol), 100 mM EDTA, 0.4 mg/ml xylene cyanol, and 0.4 mg/ml bromophenol blue]. The fraction of cleaved rSA_5_ was determined by running the reaction products on a 20% denaturing polyacrylamide gel. Measurement of this fraction for the reaction aliquots from various folding times was used to monitor the formation of the native ribozyme as a function of folding time (**47**). Substrate cleavage reactions were performed for two min, which allows for essentially complete cleavage of substrate bound to the native ribozyme and insignificant cleavage of substrate bound to misfolded ribozyme. Thus, under these conditions the fraction of the substrate that is cleaved provides a good measure of the fraction of the ribozyme that is in the native state.

## Results

### High-throughput screen uncovers pool of candidate proteins that may assist RNA folding

Given the heterogeneous nature of proteins that function as general RNA chaperones, we performed a high-throughput screen to identify proteins that affect RNA folding. We anticipated that these proteins would likely include nucleic acid-binding proteins (and especifically RBPs) with general RNA chaperone activity. For the screen, we used *in vivo* regional RNA structure probing, which provides a framework for studying RNA molecules in their cellular context. The *in vivo* RNA Structural Sensing System (iRS^3^) identifies structurally accessible regions of a transcript by detecting hybridization of 9-16 nt regions of user-designed, complementary asRNA probes (**31**). Successful binding and hybridization of the asRNA probe to a target RNA region disrupts an adjacent RBS-sequestering hairpin structure and results in translation of green fluorescence protein (GFP) (schematic shown in **Fig. 1A, bottom**). Importantly, this method has been successfully applied in different contexts to interrogate functional sites and conformational arrangements within different types of RNA molecules (**20, 21**) and to capture effects of protein interactions *in vivo* (**20**). Thus, we anticipated that this probing method could detect changes in the folding pathway of a structured target RNA upon disrupting genes encoding RNA chaperones that influence its folding. Specifically, we used the *Tetrahymena* group I intron ribozyme (herein referred to as ribozyme), since the structure and folding of this molecule have been extensively characterized by different *in vivo* and *in vitro* methods in the past (**18, 22–24**). The ribozyme has a compact native structure, and it tends to misfold into a well-defined, native-like conformation in the absence of RNA chaperones (**22**). The native and misfolded structures of the ribozyme have recently been solved using cryo-EM, providing a high-resolution view of structural differences to guide our regional accessibility profiling (**25–27**).

**Figure 1.**
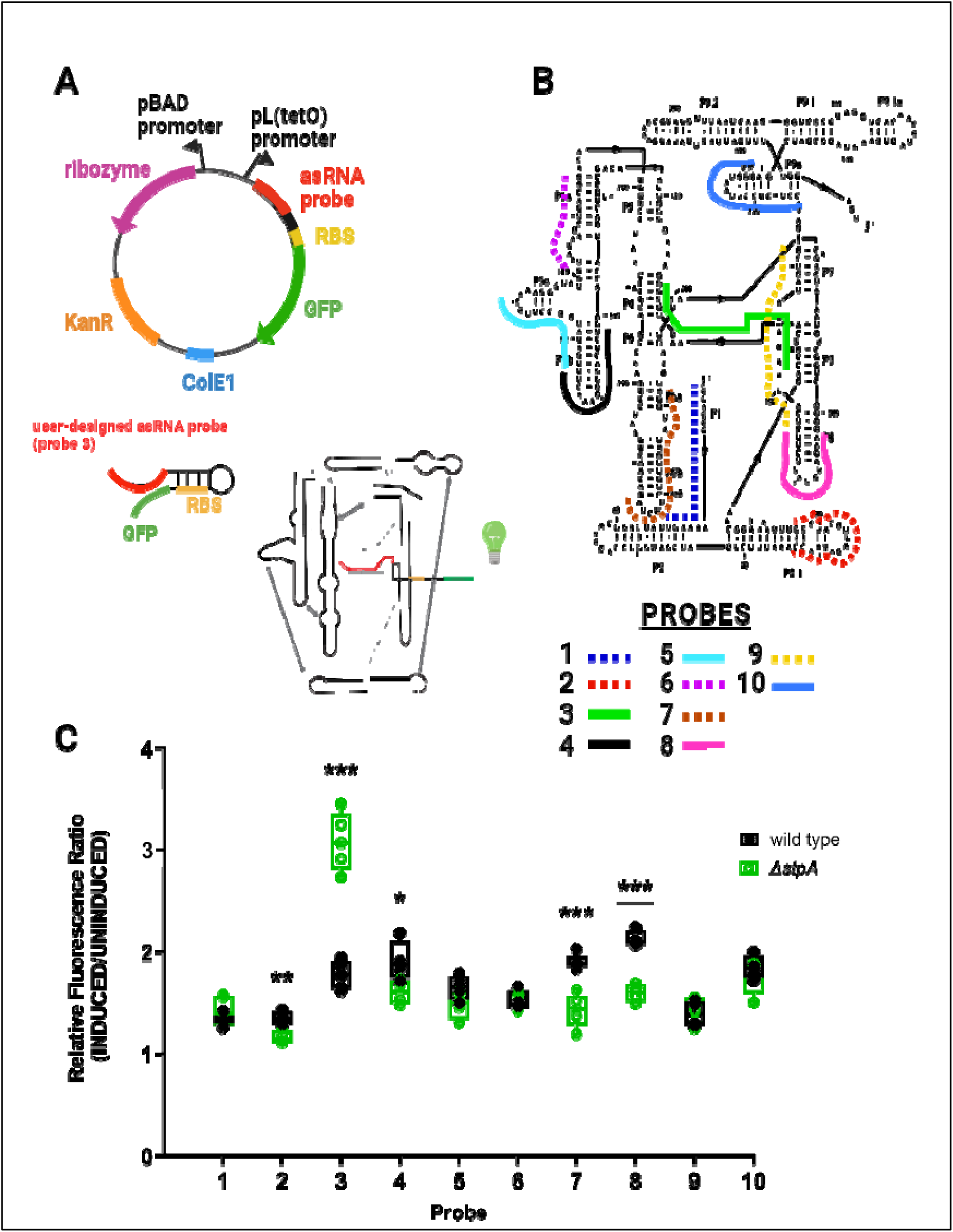
Accessibility mapping captures structural rearrangements of the *Tetrahymena* group I intron ribozyme. (A) Plasmid map and schematic representation of asRNA probing methodology used to target the ribozyme. The iRS^3^ reporter plasmid contains a pBAD promoter followed by the group I intron ribozyme sequence and GFP fused to a user-designed asRNA probe under a TetR-regulated, pL(tetO) promoter. In the iRS^3^ approach, asRNA binding alleviates sequestration of the ribosome binding site (RBS) region and allows translation of the GFP reporter. (B) Schematic of the target regions for each asRNA, designed previously (**18**) to probe the regional accessibility of the ribozyme. Probe 3 (shown in green) is the asRNA probe complementary to both the P3 and P4 domains of the ribozyme. (C) iRS^3^ fluorescence shifts corresponding to 10 probes targeting unique regions within the ribozyme expressed in *E. coli* wild type BW25113 and Δ*stpA* strains. For each probe, fluorescence ratios were calculated by dividing induced (asRNA probe + ribozyme) by target-uninduced (asRNA probe only) median fluorescence values. Relative fluorescence ratios for each probe are shown in the graph and represent the measured fluorescence for at least 4 independent biological replicates. Asterisks denote statistically significant differences between the relative fluorescence of the iRS^3^ system when expressed in the Δ*stpA* strain relative to the wild type parent strain (unpaired t-test; * p-value <0.05, ** p-value <0.01, *** p-value <0.001).

We first profiled changes in regional accessibility of the ribozyme upon deletion of the known RNA chaperone StpA, which has been shown to promote annealing and strand displacement of model RNA substrates *in vitro* (**10**). We used 10 sequence-specific asRNA probes (**18**) that target different regions of the ribozyme to profile changes in regional accessibility that could indicate chaperone-mediated structural rearrangements. Two additional control probes representative of the high and low ends of fluorescence were used in every experiment to validate probe induction and fluorescence detection (**Supplementary Fig. S4**) (**19**). The fluorescence signal for the 10 asRNA probes was measured when expressed in wildtype *E. coli* and compared to the signal measured in a single-deletion mutant *E. coli* Δ*stpA* (obtained and verified by genomic PCRs from the Keio collection, **49**). The specific regions of the ribozyme targeted by each probe are shown in **Fig. 1B**. For each probe, we compared the fluorescence ratio for the Δ*stpA* strain [the fluorescence value upon induction of the ribozyme relative to an uninduced (probe-only) control] with the analogous ratio for the wild type strain (**Fig. 1C**). Notably, upon deletion of *stpA* we observed a roughly 75% increase in fluorescence for Probe 3 (p-value < 0.001) which targets the P3 and P4 domains of the ribozyme, indicating an StpA-related change in accessibility within this region. These results agree with our previously published data showing that Probe 3 can capture accessibility changes in a well-characterized ribozyme variant that lacks important tertiary contacts and increases solvent exposure of the core (**18, 23**). The increase in Probe 3 accessibility in the absence of StpA may result from an increased population of folded conformation(s) with exposure of the complementary nucleotides and/or increased population of less stable conformations, with transient accessibility occurring during local unfolding events. This region of the ribozyme, which includes the P3 helix, displays increased exposure to footprinting reagents in the known misfolded conformation, and P3 is required to unwind and rewind during refolding from this misfolded conformation to the native state. (**22, 70**). Additionally, another well-studied chaperone, CYT-19, accelerates this transition *in vitro* and therefore reduces exposure of this region (**71**). We also observed minor, but significant, drops in the fluorescence signal of Probes 7 and 8 (∼ 25% reductions; p-value <0.001) which target primarily domains P6b and P8 respectively. These helices are on the surface of the native ribozyme structure and may be decreased in accessibility in some non-native conformations. Similarly, more subtle changes (<15% drops, p-value <0.05) at the L2.1 and L5b loops, targeted by Probe 2 and Probe 4 respectively, could be explained by changes in the accessibility of these regions. Importantly, confirmation of the ability to capture established RNA conformational changes upon deletion of the known RNA chaperone StpA served as a proof of concept for a larger screen, motivating the use of this accessibility probing approach to screen for additional general chaperone proteins that contribute to the folding of the ribozyme.

Thus, we designed a Tn5-transposon library and incorporated the iRS^3^ system expressing Probe 3 to screen for genes encoding candidate RNA-chaperone proteins. The library was prepared to ensure coverage of 3x the number of suspected genes in *E. coli* (see **Materials and Methods**). As illustrated in **Fig. 2**, transposed strains harboring the reporter Probe 3 plasmid were sorted based on their higher levels of fluorescence relative to wild type *E. coli* using fluorescence-activated cell sorting (FACS). Like when probing in the presence and absence of the StpA chaperone (albeit we did not detect StpA in our screen, see **Discussion** for possible reasons), we attributed high Probe 3 fluorescence in transposed strains to changes in the folding dynamics of the ribozyme when expressed *in vivo* due to disrupting a gene that influences RNA folding. After fluorescence sorting, the isolated mutants were subjected to a second screen to validate their fluorescence signal (**Supplementary Fig. S1**, log2 fold-change cutoff of >0.5). From this second screen, high-fluorescence mutants were pooled, and transposon insertions were identified through whole-genome sequencing (WGS) (see **Materials and Methods)**. From this pool, 31 unique transposon insertions were mapped within different genes (listed in **Table 1**) and were identified as potential RNA chaperone candidates. Interestingly, we did not find significant enrichment of proteins in any specific pathways, supporting the view that general RNA chaperone proteins may have distinct primary functions and ‘moonlight’ as chaperones (**4**). The identified candidate genes encode proteins that participate in a wide variety of cellular processes including cAMP biosynthesis, transmembrane transport, plasmid recombination, amino acid metabolism, and response to metal ions, among others.

**Figure 2.**
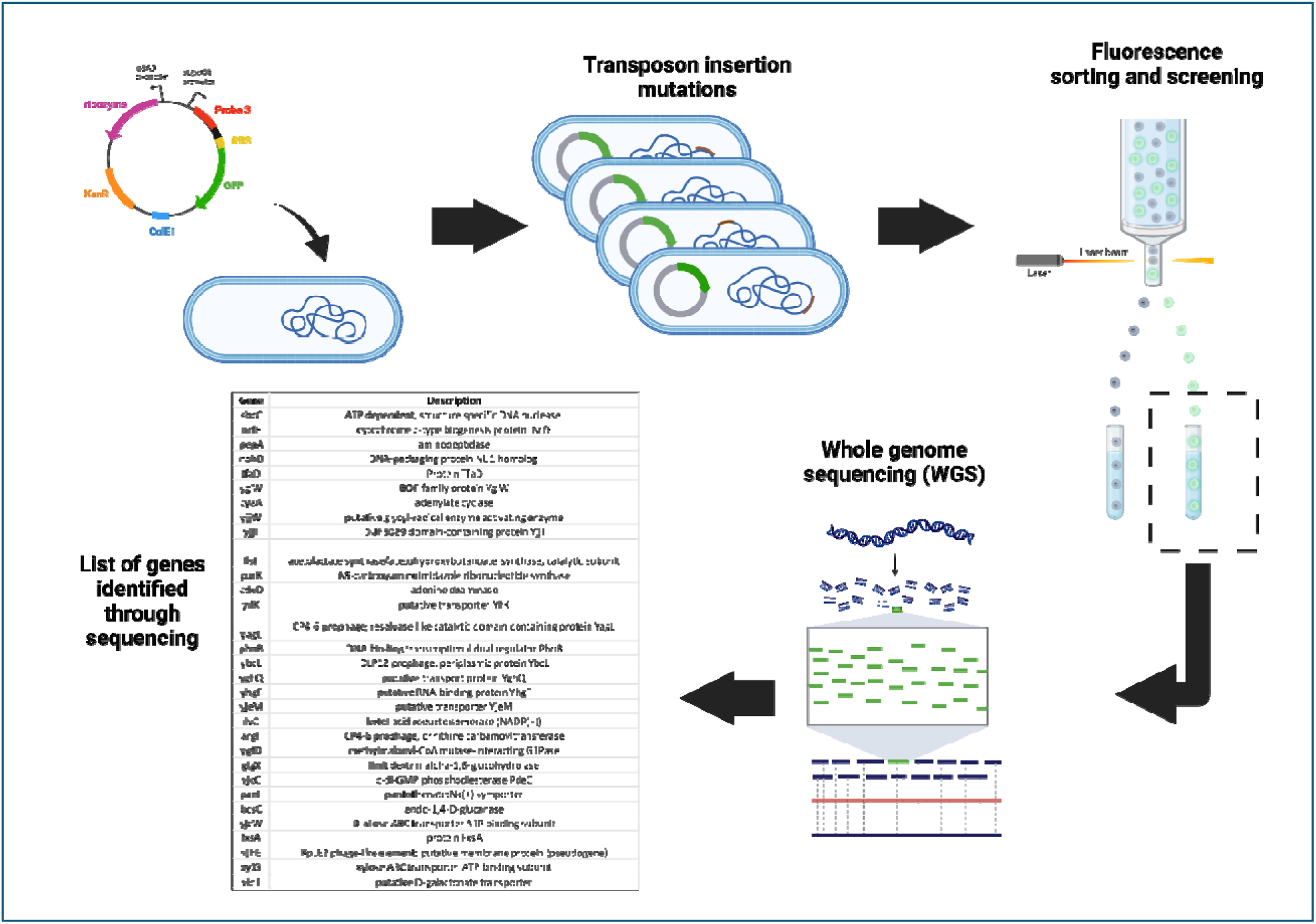
Methodology to uncover candidate proteins that affect RNA folding in *vivo*. A library of *E. coli* MG1655 cells with random transposon insertions of the TetR cassette was generated by adapting previously published procedures (**77**). The plasmid containing the iRS^3^ system with the Probe 3 asRNA was transformed into the library. Cells were sorted based on their GFP expression. High-fluorescence isolates were prepped for whole-genome sequencing, allowing for transposon insertion mapping. Image created with BioRender.com.

### RNA accessibility profiling of the Tetrahymena group I intron ribozyme in single-gene knockout strains identifies PepA and YagL as putative RNA chaperones

To further validate candidate genes and rule out multiple transposon insertions in our pooled strains, we performed further accessibility measurements on single-gene knockouts of our candidate chaperone-encoding genes. Specifically, we sought to validate changes in Probe 3 fluorescence (reflecting changes in the regional accessibility at the P3/P4 region of the ribozyme) by transforming the iRS^3^ reporter carrying Probe 3 into single-gene knockout strains of the identified candidate genes **(Table 1)**. To conduct these experiments, we cured out the KanR antibiotic marker from single-gene knockout strains in the Keio collection for 16/31 of our candidate genes using FLP recombination and confirmed the removal by colony PCR (**Supplementary Table S1**). Removing the antibiotic marker ensured that the strains were compatible with the accessibility reporter plasmid, which contains a kanamycin resistance cassette, and reduced the potential metabolic burden of using multiple antibiotics. Of the remaining 15 candidate genes, we were not able to obtain knockout strains for 10 of them (*nrfE, nohD, tfaD, yjjW, purK, adeD, ygfD, yjcW, xylG,* and *yidT*) and we were not successful at curing out the KanR cassette for five Keio collection strains (*yjjI, ilvC, glgX, yjcC* and *panF*).

We validated the expected fluorescence increases, indicating enhanced accessibility for the region targeted by Probe 3, for 8/16 tested strains relative to the wild type strain (*sbcC, pepA, yglW, yagL, argF,yifK, fxsA* and *yjhE*) (p-value <0.05; **Fig. 3**). Of these eight genes, we were particularly intrigued by *pepA* and *yagL*. The aminopeptidase A protein, PepA, was previously shown to be a multifunctional protein with DNA-binding capabilities (**50, 51**). Similarly, the *yagL* gene was predicted to encode a DNA-binding recombinase. Because previously characterized general RNA chaperones in *E. coli* (CspA, StpA) are DNA-binding proteins that moonlight as RNA remodelers, we pursued investigations of the roles of PepA and YagL in RNA folding. We excluded SbcCD subunit C (SbcC) from further studies because *in vitro* reconstitution of this complex for biochemical confirmation studies presented a challenge given that this protein forms a complex with subunit D (SbcD) for its DNA nuclease function (**52**). Additionally, we chose not to prioritize the uncharacterized genes *yifK, fxsA* and *yjhE* for further investigation.

**Figure 3.**
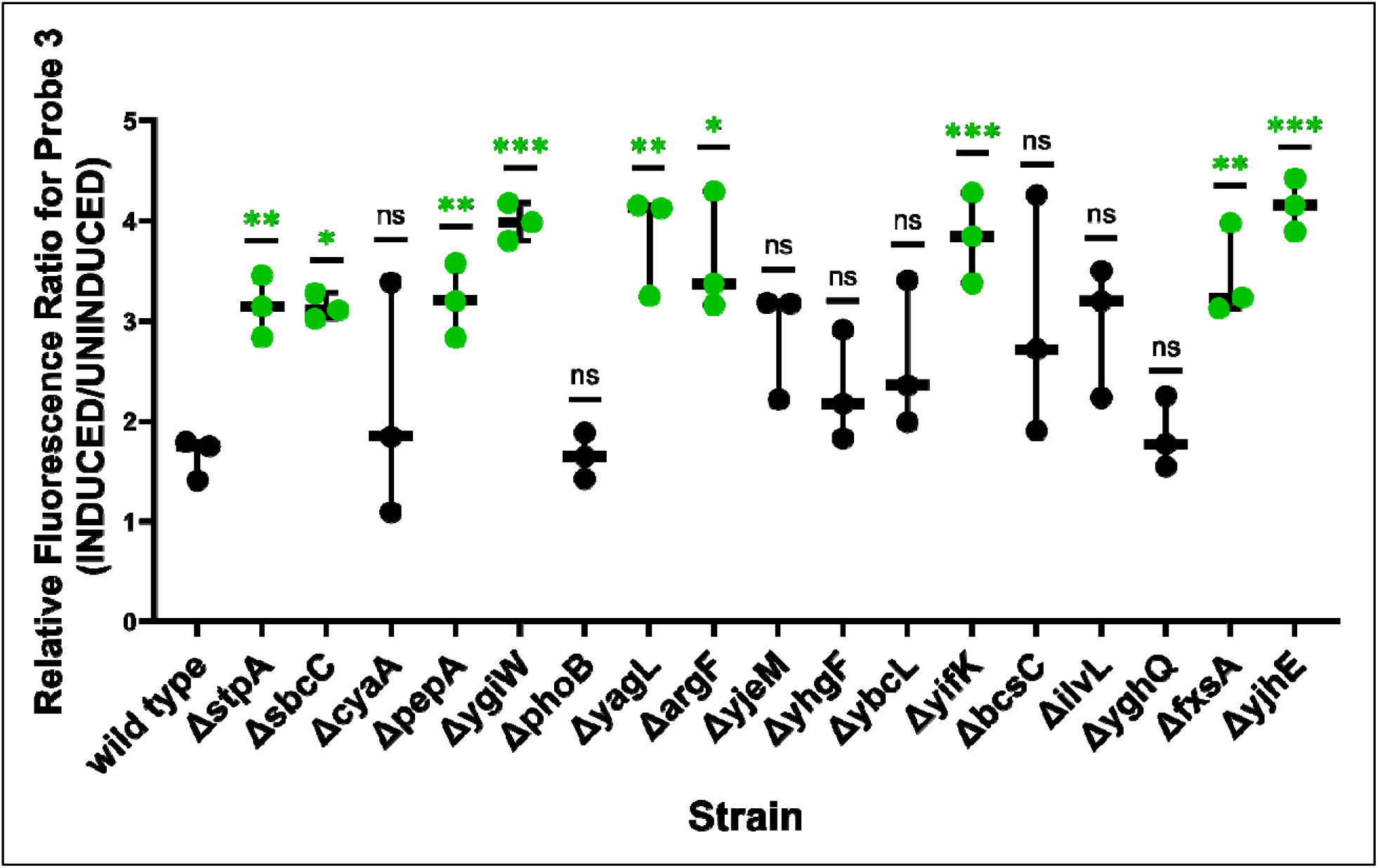
Fluorescence of iRS^3^-targeting of ribozyme region P3/P4 on single-gene knockouts validated 8/16 cellular factors. Fluorescence ratios were calculated by dividing induced (Probe 3 asRNA + ribozyme) by target-uninduced (Probe 3 asRNA only) median fluorescence values. Individual data points shown in the graph represent the paired induced/uninduced ratio obtained for three independent biological replicates. Asterisks denote significant difference between the fluorescence ratio of Probe 3 when expressed on the single-knockout strain relative to when expressed on wildtype *E. coli* BW25113 (unpaired t-test; * p-value <0.05, ** p-value <0.01, *** p-value <0.001). *E. coli* Δ*stpA* was included as a positive control for these experiments.

To better understand the influence of PepA and YagL on the folding of the ribozyme, we evaluated the accessibility of additional regions using the full set of asRNA probes for this target [(**18, 53**); targeted regions are listed in **Table 2**]. As shown in **Fig. 4A**, when using additional ribozyme-specific asRNA probes on a Δ*pepA* strain expressing the *Tetrahymena* ribozyme, we observe a similar fluorescence pattern to that generated on the Δ*stpA* strain (results shown in **Fig. 1).** Specifically, Probe 3 (targeting the P3/P4 domain) shows significantly higher fluorescence in the absence of the PepA protein (∼ 150% increase, p-value <0.001). We also observed modest but significant drops in the fluorescence of Probe 7 and Probe 8 [∼ 20 % (p-value < 0.05) and ∼40 % (p-value < 0.01) reductions] which target the P6b helix and the adjacent tetraloop-receptor (J6a/6b) and the L8 loop, respectively. A minor increase in fluorescence was also detected for the regions targeted by Probe 6 (∼30% increase, p-value <0.001). Because the fluorescence changes are not uniform for the different probes, we infer that the increase in fluorescence observed when using Probe 3 is due at least in part to structural changes that result in increased accessibility at the region targeted by this probe. This was further validated via Northern blotting by probing transcript levels of each asRNA using a GFP-specific probe, which ruled out intracellular probe concentration as a contributing factor to the observed changes in fluorescence (**Supplementary Fig. S5**).

**Figure 4.**
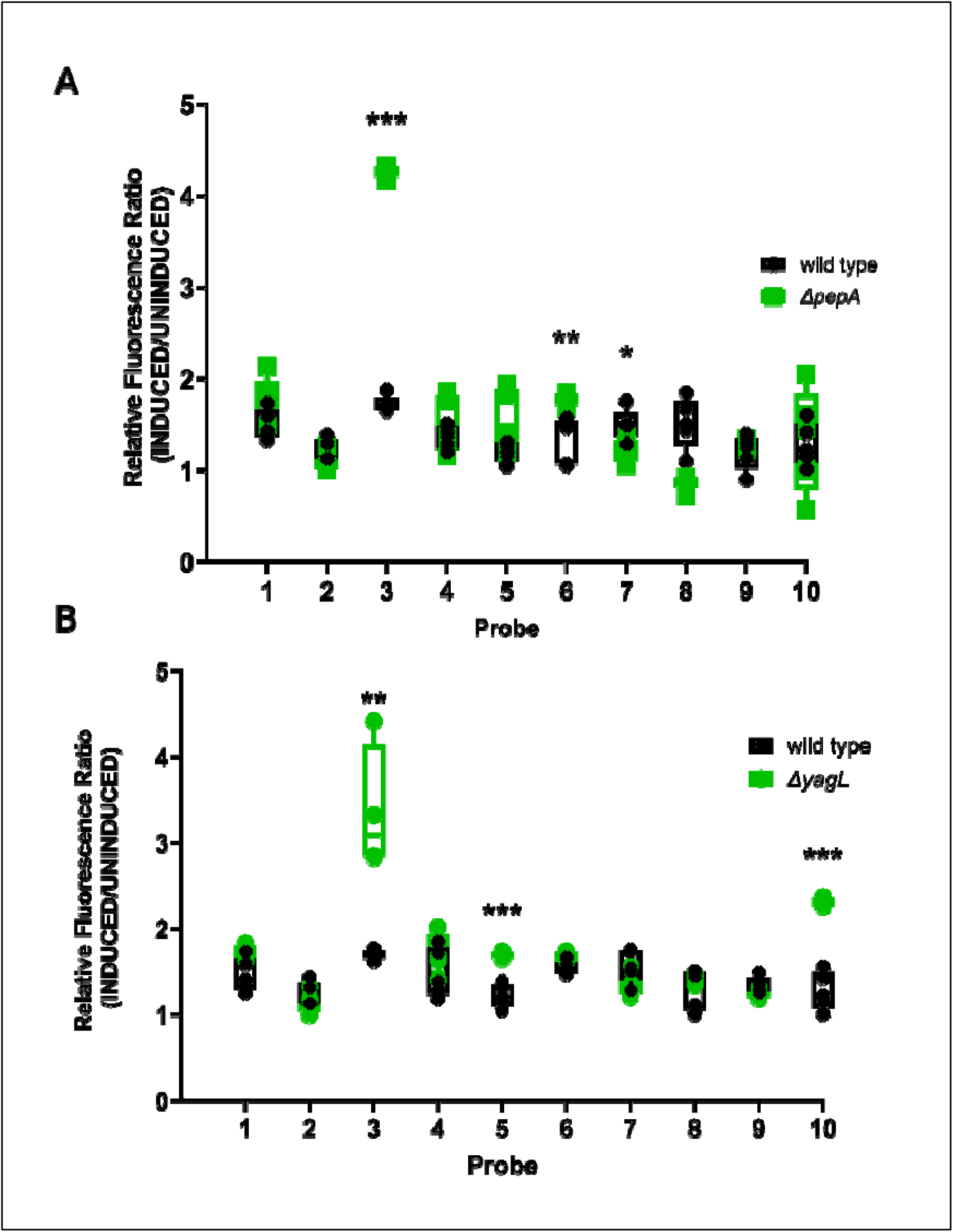
Protein-mediated changes on the accessibility profile of the *Tetrahymena* ribozyme suggest RNA chaperone role for PepA and YagL. (A) Fluorescence of iRS^3^- targeting of the *Tetrahymena* ribozyme when expressed in *E. coli* wild type BW25113 and Δ*pepA*. For each probe, fluorescence ratios were calculated by dividing paired induced (asRNA probe + ribozyme) by target-uninduced (asRNA probe only) median fluorescence values. Individual data points shown in the graph (‘□’. Δ*pepA* and ‘x’ -wild type) represent the obtained values for independent biological quadruplets. (B) Ribozyme accessibility profile captured by the iRS^3^ assay when expressed in *E. coli* wild type and Δ*yagL.* For each probe, fluorescence ratios were calculated by dividing induced (asRNA probe + ribozyme) by uninduced (asRNA probe only) median fluorescence values for independent biological quadruplets (depicted as ‘○’-Δ*yagL* and ‘x’ -wild type in the graph). Asterisks denote significant difference in the fluorescence ratio for observed in the mutant strain (Δ*pepA* or Δ*yagL)* relative to that of the wild type strain (unpaired t-test; * p-value <0.05, ** p-value <0.01, *** p-value <0.001).

**Table 2.**
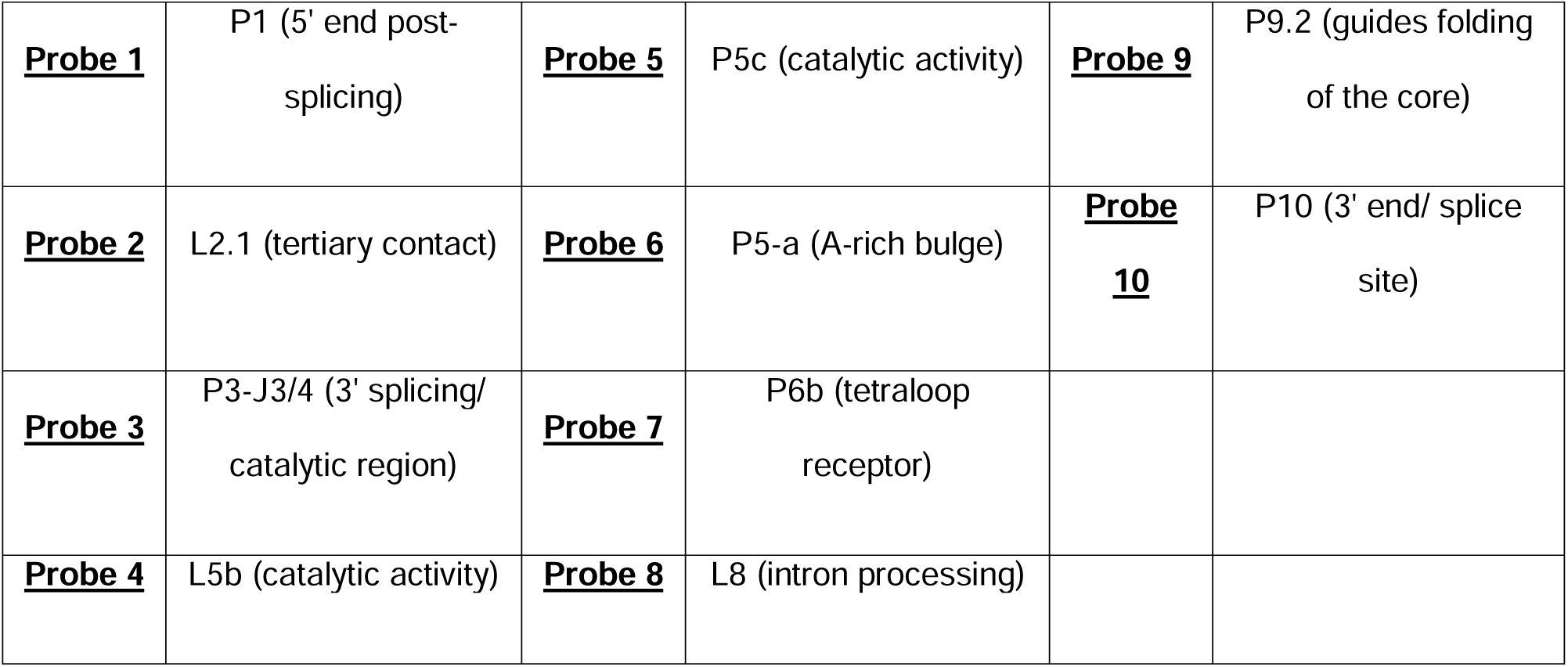
Regions targeted by the 10 asRNA probes used to profile the *Tetrahymena* gI intron ribozyme. Ten asRNA probes previously published in Sowa et al. (2015) (**18**) were used to measure changes in regional accessibility for specific regions along the ribozyme transcript.

|  |  |  |  |  |  |
| --- | --- | --- | --- | --- | --- |
| **<u>Probe 1</u>** | P1 (5' end post-splicing) | **<u>Probe 5</u>** | P5c (catalytic activity) | **<u>Probe 9</u>** | P9.2 (guides folding of the core) |
| **<u>Probe 2</u>** | L2.1 (tertiary contact) | **<u>Probe 6</u>** | P5-a (A-rich bulge) | **<u>Probe 10</u>** | P10 (3' end/ splice site) |
| **<u>Probe 3</u>** | P3-J3/4 (3' splicing/ catalytic region) | **<u>Probe 7</u>** | P6b (tetraloop receptor) |  |  |
| **<u>Probe 4</u>** | L5b (catalytic activity) | **<u>Probe 8</u>** | L8 (intron processing) |  |  |

When we measured the fluorescence of the ribozyme-specific asRNA probes on the Δ*yagL* strain expressing the ribozyme (results shown in **Fig. 4B**), we obtained a similar fluorescence change for Probe 3 (targeting the P3 and P4 domains) to that observed for the Δ*stpA* and Δ*pepA* strains, relative to the wildtype strain (∼98% increase, p-value <0.01). Additionally, we observed unique changes in fluorescence, relative to the wildtype strain, for Probe 5 and Probe 10 (targeting the P5c hairpin and the 3’ end/splice site region, respectively). For Probe 5, we observed a ∼ 38% increase in GFP reporter fluorescence (p-value <0.001). This probe targets P5c, part of the P5abc domain, which stabilizes the catalytic core of the ribozyme by forming tertiary contacts and forms rapidly during folding of the ribozyme (**54**). Similarly, Probe 10 shows an ∼ 82% increase in fluorescence (p-value <0.001). Probe 10 targets the 3’ end and the P9 domain of the ribozyme, which participate in peripheral interactions and may be increased in accessibility in the misfolded conformation and during refolding to the native state (**25–27**). We also confirmed that fluorescence shifts are not due to differences in transcript levels of the accessibility probes across strains via Northern blotting analysis using a GFP-specific radiolabeled probes **(Supplementary Fig. S5)**. We note that the Northern blots showed a significant reduction in transcript levels for the iRS^3^ probe in the Δ*yagL* strain. However, we concluded that the observed increases in fluorescence signal from the accesibility probing assay are likely due to changes in the abilities of the probes to hybridize to their target regions; since a reduction in probe levels would only reduce the magnitude of the observed changes, and no significant reductions in fluorescence were observed for any of the probed regions.

Together, these results isolated PepA and YagL as two proteins that are likely capable of remodeling RNA *in vivo*, as determined by changes in accessibility patterns of key regions within the ribozyme upon their deletion. We did not perform further tests on other genes identified via the high throughput screen, but those genes represent promising candidates for future work.

### YagL accelerates native refolding of the Tetrahymena group I intron ribozyme in vitro from a misfolded state

To investigate how PepA and YagL affect folding of the ribozyme, we tested the ability of these proteins to accelerate native refolding from the long-lived misfolded conformation. To measure refolding, we used a two-step, or ‘discontinuous’, folding assay in which catalytic activity is used to monitor native ribozyme folding **(Fig. 5A).**In this experiment, the ribozyme is pre-incubated with Mg^2+^ to generate the misfolded conformation. Then, during the first assay step, the ribozyme is allowed to refold to the native state in the presence of various concentrations of chaperone protein (or no chaperone protein as a control), with reaction aliquots stopped and removed at various times. In the second step, the fraction of native ribozyme is determined for each of these time points by measuring the fraction of an added oligonucleotide substrate that is rapidly cleaved by the native ribozyme (**22, 55**) **44**).

**Figure 5.**
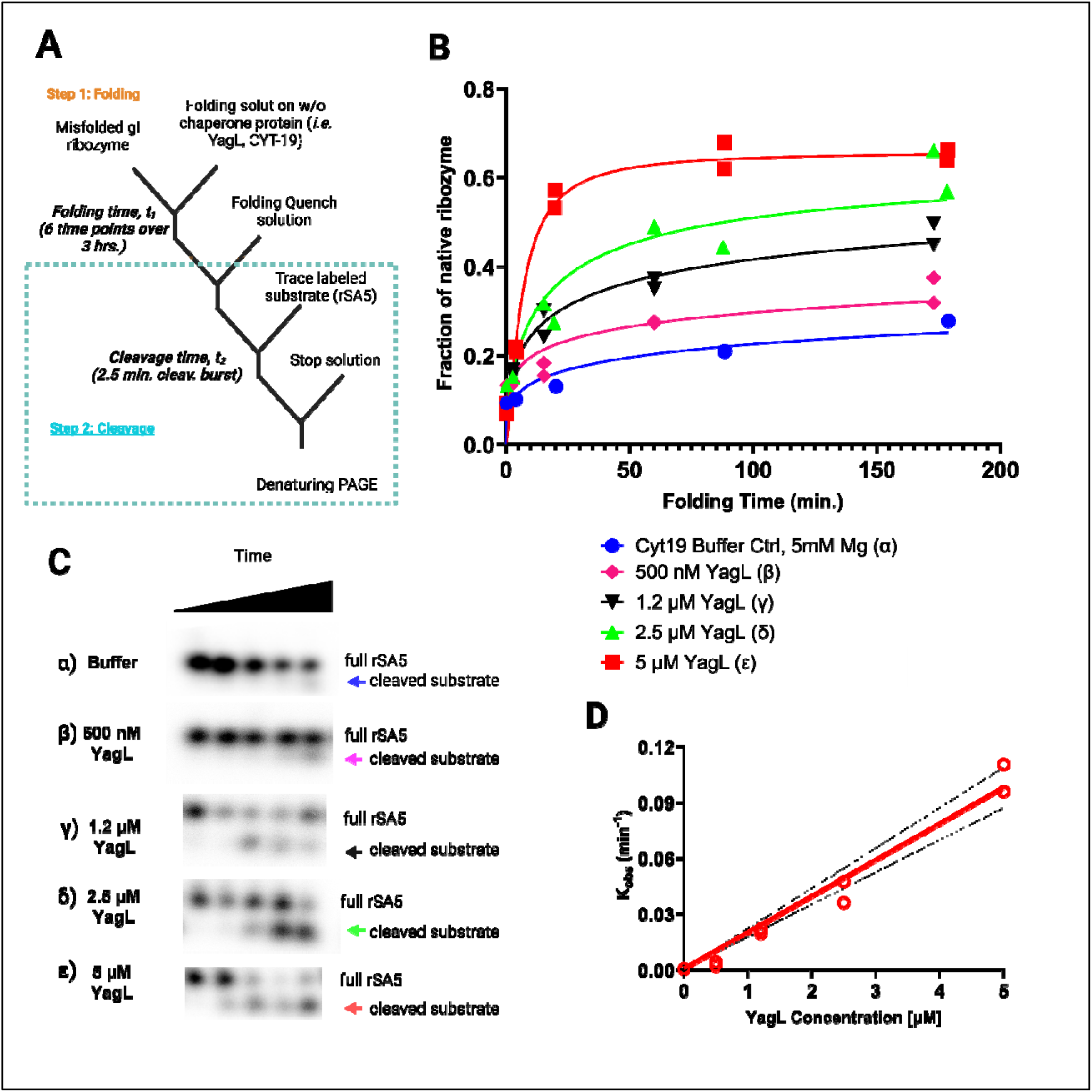
Acceleration of native ribozyme folding by YagL. (A) The *Tetrahymena* gI intron ribozyme was pre-incubated into a misfolded state and then added into folding reactions without YagL (α), with 500 nM YagL (β), 1.2 µM YagL (γ), 2.5 µM YagL (δ), or 5 µM YagL (ε). Reactions were stopped at different times, after which radiolabeled rSA5 (which mimics the 5’-splice-site junction cleaved by the ribozyme) was added to perform substrate cleavage reactions. After quenching, reactions were stopped and analyzed by denaturing PAGE to quantify product formation. (B) The fraction of cleaved substrate was quantified and used to generate plots of the fraction of native ribozyme as a function of folding time. From these plots, average rate constants from two independent determinations of native state formation were 0.0008 min^-1^ (α), 0.0032 min^-1^ (β), 0.0210 min^-1^ (γ), 0.0421 min^-1^ (δ), and 0.100 min^-1^ (ε). The faster reactions gave end points of 0.66, and end points for the slower reactions were forced to this value. (C) Gel images used to quantify cleavage product formation (these images were obtained for one out of the two independent determinations performed; raw and processed images for all replicates can be found in the **Supplementary Information- Appendix 3** for this publication). The cleaved substrate, indicated by the arrows, increases with folding time and with YagL concentration. (D) Observed rate constants from (B) plotted against YagL concentration. Dotted black lines denote the 95% confidence intervals. From this linear fitting, the second-order rate constant K_Cat_/K_M_ was determined to be ∼1.9 x 10^4^M^-1^min^-1^ in the presence of 5 mM Mg^2+^.

Using this assay, we found that purified YagL from *E. coli* accelerates native refolding of the ribozyme in a concentration-dependent manner **(Fig. 5B-C)**. In the absence of YagL, refolding occurred slowly, on the time scale of hours, consistent with previous work (**22, 55**). We plotted the observed rate constants against YagL concentration, obtaining a k_Cat_/K_M_ value of 1.9 + 0.1 x 10^4^ M^-1^ min^-1^ **(Fig. 5D)**. This value is comparable to that for the established CYT-19 RNA chaperone under similar experimental conditions (**56**). The data also revealed the possibility of upward curvature in the concentration dependence, which might indicate the cooperative involvement of multiple protein molecules. Importantly, in this assay YagL was inactivated by proteinase K after the folding step and before the substrate cleavage step. Thus, the experiment demonstrates that YagL functions as a chaperone in this folding reaction, via increasing the rate constant for ribozyme refolding to the native state, not via direct modulation of the catalytic activity of the ribozyme.

In contrast, purified PepA did not accelerate ribozyme refolding compared to the control reaction lacking PepA **(Supplementary Fig. S6)**. We did not evaluate whether PepA plays other chaperone roles such as destabilization of local structures within the ribozyme. Future work should further investigate the functional role and relevant conditions in which PepA interacts productively with RNA substrates.

### PepA and YagL are RNA-binding proteins that bind single-stranded and double-stranded RNA

Previously characterized general chaperones in *E. coli* (such as CspA and StpA) bind RNA with modest affinities in the µM range. Further, mutations that increase the RNA-binding affinity of StpA resulted in reduced chaperone activity (as measured by the ability of StpA to promote cis-splicing of the *td* intron) (**10**), suggesting that there may be an optimal affinity range for RNA chaperone activity.

Although DNA-binding activity has been shown for PepA (**28, 57**) and predicted for YagL (**58**), their ability to directly bind RNA was unknown. Therefore, we used nitrocellulose filter binding to measure binding of these proteins to three 5’-^32^P-labeled short RNA oligonucleotides that were used previously to evaluate StpA-RNA binding (**10**). Specifically, we used two random 21-mer RNA sequences (“21R+” and its complementary sequence “21R-“) to evaluate binding to ssRNA, and we used a short hairpin loop (herein referred to as “hairpin” for simplicity) to evaluate binding to a dsRNA. As shown in **Fig. 6A-C**, PepA bound both 21R+ and 21R-with affinities in the low µM range (2.0 ± 0.6 µM and 0.6 ± 0.3 µM, respectively), demonstrating that this protein is capable of binding ssRNA. Further, PepA bound to the hairpin oligonucleotide with an estimated *K*_D_ value of 0.6 ± 0.4 µM, suggesting that it can bind both ssRNA and dsRNA without an apparent substrate preference. Likewise, YagL bound RNA in the low μM range, with a detectable preference for dsRNA. YagL bound both 21R+ and 21R-with similar affinities (4.5 ± 0.3 µM and 5.0 ± 1.2 µM, respectively), and it bound the hairpin 2-fold more tightly (2.4 ± 0.3 µM; **Fig. 7C**). For the ssRNAs, the binding curves displayed hints of sigmoidal behavior, raising the possibility of cooperative binding, but the fitted Hill coefficients were close to one, and for simplicity we fit these curves using a simple, hyperbolic binding equation (**Fig. 7C**).

**Figure 6.**
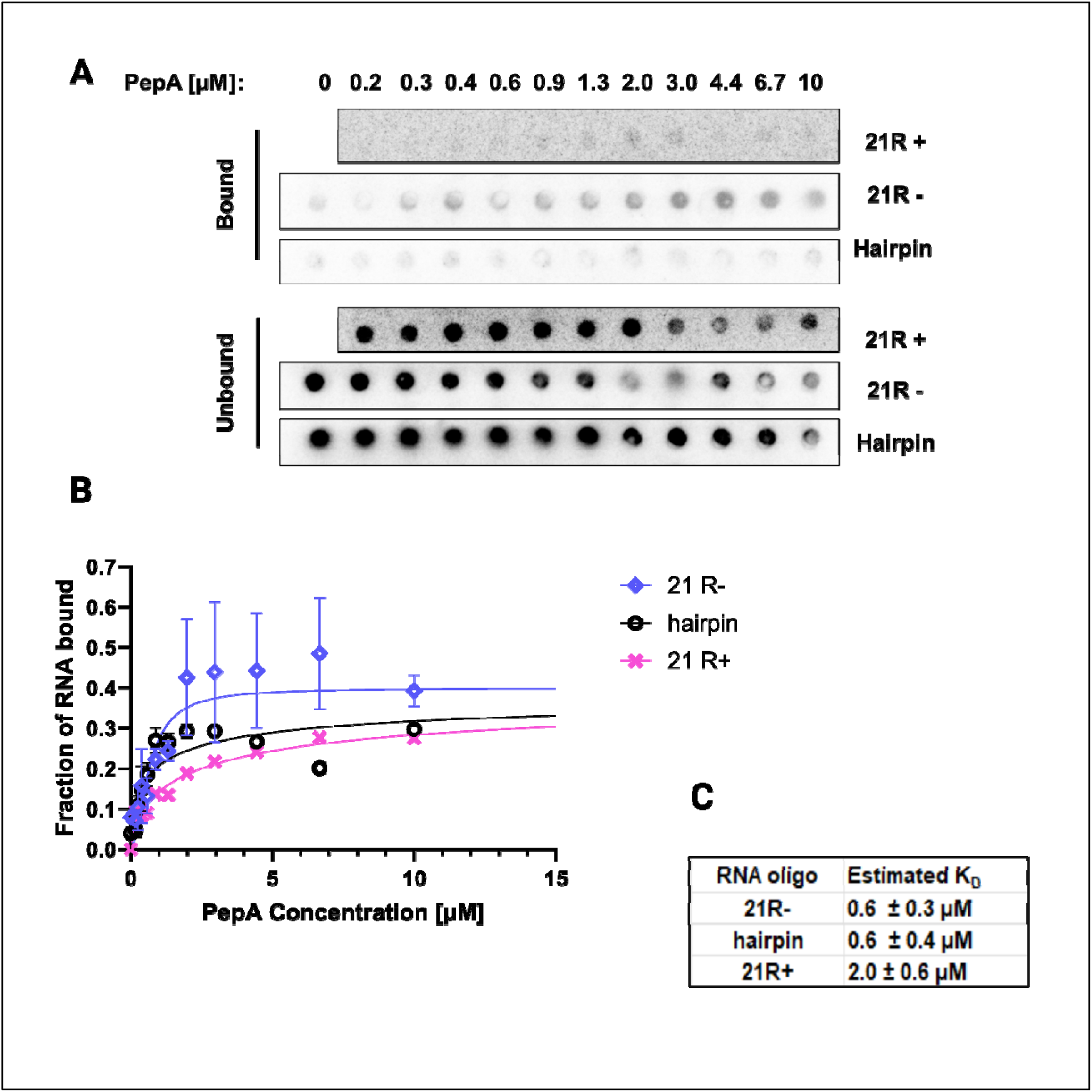
PepA binds both ssRNA and dsRNA. Filter binding assays were performed to test binding of PepA to two single-stranded 21mer RNA sequences: “21R+” (*5*′*-AUGUGGAAAAUCUCUAGCAGU-3*′) and “21R-” (*5*′*-CUGCUAGAGAUUUUCCACAU-3*′). A short hairpin loop oligo (*5’-GCTCTAGAGCATTATGTTCAGATAAGG-3’*) was used to evaluate binding to small structured RNAs. (A) Representative membrane images of the bound and unbound signals. Filter binding experiments were conducted in experimental duplicates for each reaction. (B) Non-linear fits were used to generate binding curves for PepA and the three RNA substrates (“21+”, “21-“, and “hairpin”). Error bars represent the variation between the experimental replicates. (C) Estimated *K*_D_ values for each substrate.

**Figure 7.**
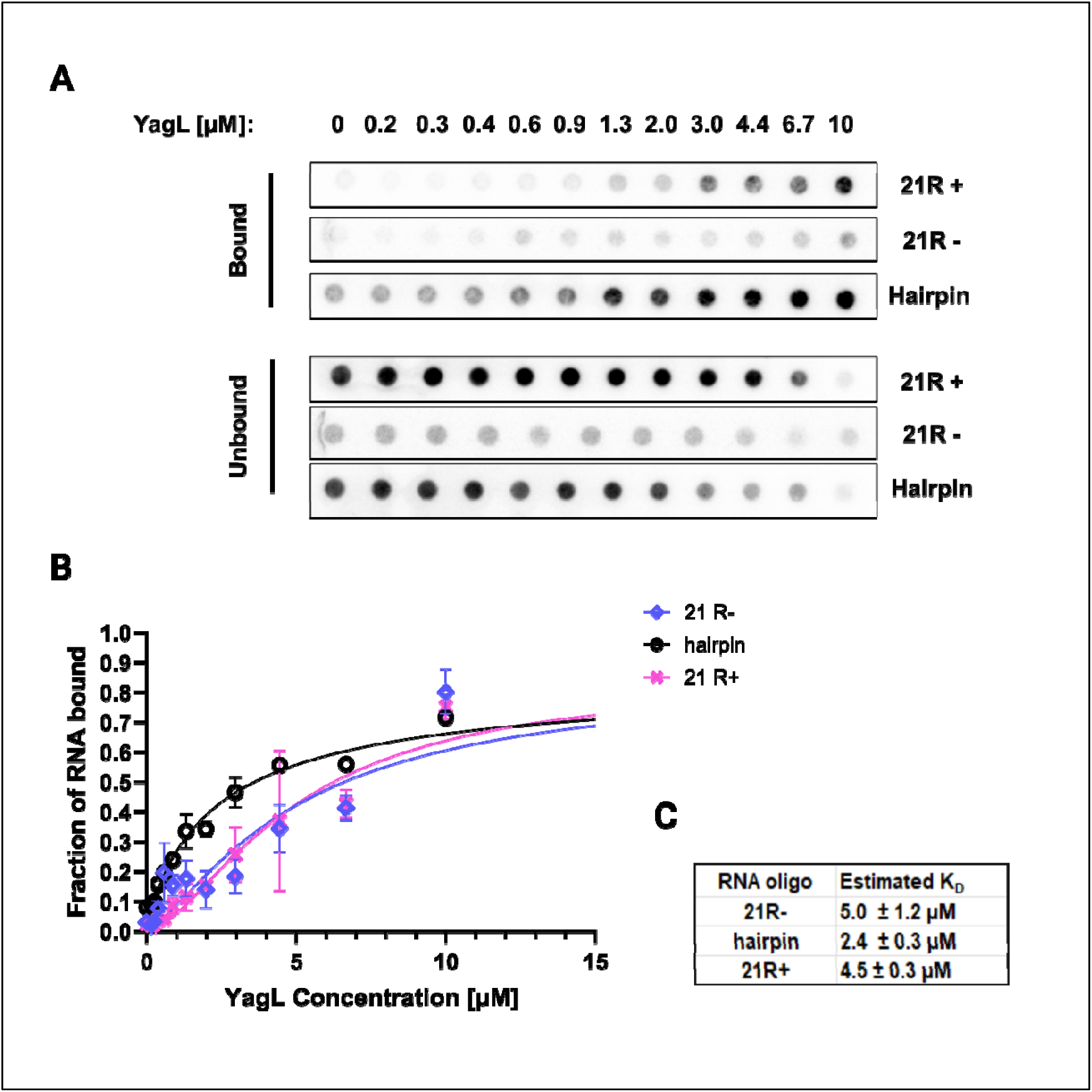
YagL binds preferentially to dsRNA. Filter binding assays were performed to test binding of YagL to two single-stranded 21mer RNA sequences: “21R+” (*5*′*-AUGUGGAAAAUCUCUAGCAGU-3*′) and “21R-” (*5*′*-CUGCUAGAGAUUUUCCACAU-3*′). A short hairpin loop oligo (*5’-GCTCTAGAGCATTATGTTCAGATAAGG-3’*) was used to evaluate binding to small structured RNAs. (A) Representative membrane images of the bound and unbound signals. Filter binding experiments were conducted in experimental duplicates for each reaction. (B) Non-linear fits were used to generate binding curves for YagL and the three RNA substrates (“21+”, “21-“, and “hairpin”). Error bars represent the variation between the experimental replicates. (C) Estimated *K*_D_ values for each substrate.

These results suggest that PepA and YagL modestly interact with RNA (µM range), like the general *E. coli* RNA chaperones StpA and CspA. For contrast, StpA was shown to bind short ssRNA with a *K*_D_ value of ∼580 nM in filter binding assays and to bind structured RNAs with lower affinity (ranging from 12.3-24.7 μM depending on the RNA substrate) in isothermal titration calorimetry (ITC) measurements (**10, 59**). Similarly, CspA has been reported to interact with its natural partner ACB (anti-cold box) with a *K*_D_ value of ∼12 μM (**8**).

### A structurally conserved helix-turn-helix (HTH) domain mediates YagL interactions with nucleic acids

While PepA is a moonlighting protein whose multiple functions have been documented in the literature (**28, 60**) and whose crystal structure has been solved (**61**), little was known about the structure and function of YagL. After identifying YagL as a novel RNA chaperone in *E. coli* and confirming its ability to both accelerate refolding of the ribozyme and interact with ssRNA and dsRNA substrates *in vitro*, we wanted to further investigate the structural features of YagL that could explain its RNA chaperone role.

YagL was an uncharacterized, 27.2 kDa protein predicted to function as a DNA recombinase (**58**). To determine an initial protein structural model for YagL and to identify structurally similar proteins, we subjected its full sequence to homology modeling via Phyre2. Using this approach, YagL shows structural similarity to resolvase proteins **(Supplementary Information-Appendix 1).** In parallel, we used deep learning protein structure predictions to generate a model for YagL via AlphaFold (**62**). Notably, this approach (in addition to homology modeling predictions via Phyre2), led to the identification of a helix-turn-helix (HTH) domain in the C-terminus of YagL (**Supplementary Fig. S7, panels A&B**). The HTH domain found in YagL is predominantly composed of highly conserved, positively charged residues that could mediate charge-charge interactions with RNA substrates (**Supplementary Fig. S7, panels D&F**). This binding mode is supported by additional molecular docking simulations that identified residues like Lys, Arg, His, and Trp in the HTH domain of YagL as DNA-interfacing residues that could also mediate RNA binding (**Supplementary Fig. S8, Supplementary Information- Appendix 1).** In these simulations, 6 DNA ligands (**Supplementary Table S3**) collected from homologous protein structures identified in the Phyre2 analysis were used as a proxy to investigate RNA binding (as many of the amino acid residues responsible for binding these biomolecules overlap (**63–65**)).

HTH domains are becoming increasingly identified in multifunctional proteins that bind to nucleic acid substrates (*e.g.,* transcription factors and proteins of the La domain family which serve an RNA chaperone role in eukaryotes (**66**)). Thus, we decided to evaluate if this predicted HTH domain in YagL was responsible for its RNA-binding and RNA chaperone roles by creating two YagL protein truncations; YagL-dHTH, which included the N-terminal domain of the protein minus the predicted HTH domain, and YagL-HTH, which comprised the predicted C-terminal HTH domain (**Fig. 8A**, detailed sequences are included in **Supplementary Table S4**).

**Figure 8.**
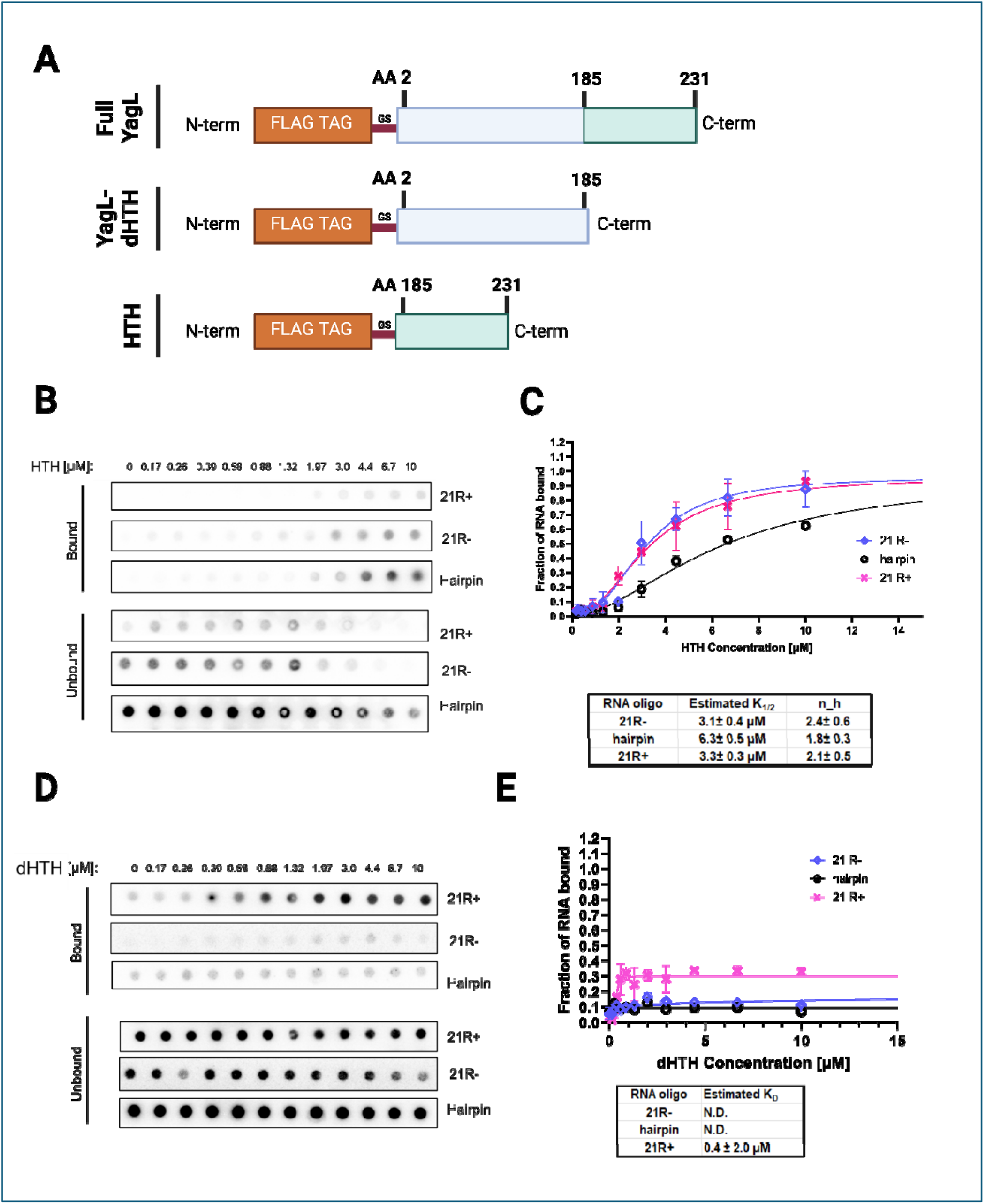
The predicted HTH domain of YagL is responsible for its RNA-binding activity. (A) Schematic visualization of the two YagL protein truncations constructed to determine their contribution to RNA-binding. YagL-dHTH includes the N-terminus domain of the protein minus a predicted HTH domain located at the C-terminus of the protein. HTH consists of the last 46 amino acids of YagL. Nucleotide sequences of these protein truncations are included in **Supplementary Table S4**. (B&D) Filter binding assays were performed to evaluate binding of the HTH and dHTH protein truncations to two single-stranded 21mer RNA sequences, “21R+” and “21R-”, and a small, structured RNA, “hairpin”. (C&E) The bound and unbound fractions were quantified to generate binding curves for the HTH and the dHTH truncated protein respectively. For the HTH truncation, sigmoidal curves were obtained, suggesting cooperative binding of ∼2 functional units (Hill coefficients: 2.4± 0.6 (21R-), 1.8± 0.3(hairpin), and 2.1± 0.5 (21R+). Image created with BioRender.com.

We performed filter binding assay experiments using the purified YagL-dHTH and YagL-HTH protein truncations. As shown in **Fig. 8 panels B&C**, the predicted HTH domain alone was able to bind RNA with estimated affinities between 3-6 µM depending on the RNA substrate used. However, unlike the full YagL protein, YagL-HTH displayed a reversed binding preference relative to the full-length YagL, with 2-fold tighter binding for the ssRNAs.

Additionally, there was pronounced upward curvature in the binding curves for all the RNAs, giving Hill coefficients of approximately two and indicating cooperative binding. For the YagL-dHTH protein, minimal oligonucleotide binding was observed at any tested protein concentration (**Fig. 8 panels D&E**), suggesting that the C-terminus region of YagL, encoding the predicted HTH domain, is responsible for most of the RNA-binding activity of YagL.

To evaluate if the predicted HTH domain was also responsible for the RNA chaperone activity of YagL, its ability to accelerate native refolding of the ribozyme was assessed using the two-step folding assay described above. As shown in **Fig. 9**, only the full YagL protein was capable of significantly accelerating the refolding of the ribozyme. Thus, we conclude that while the C-terminus HTH domain of YagL is sufficient to bind RNA substrates, the full protein is needed for its RNA chaperone function.

**Figure 9.**
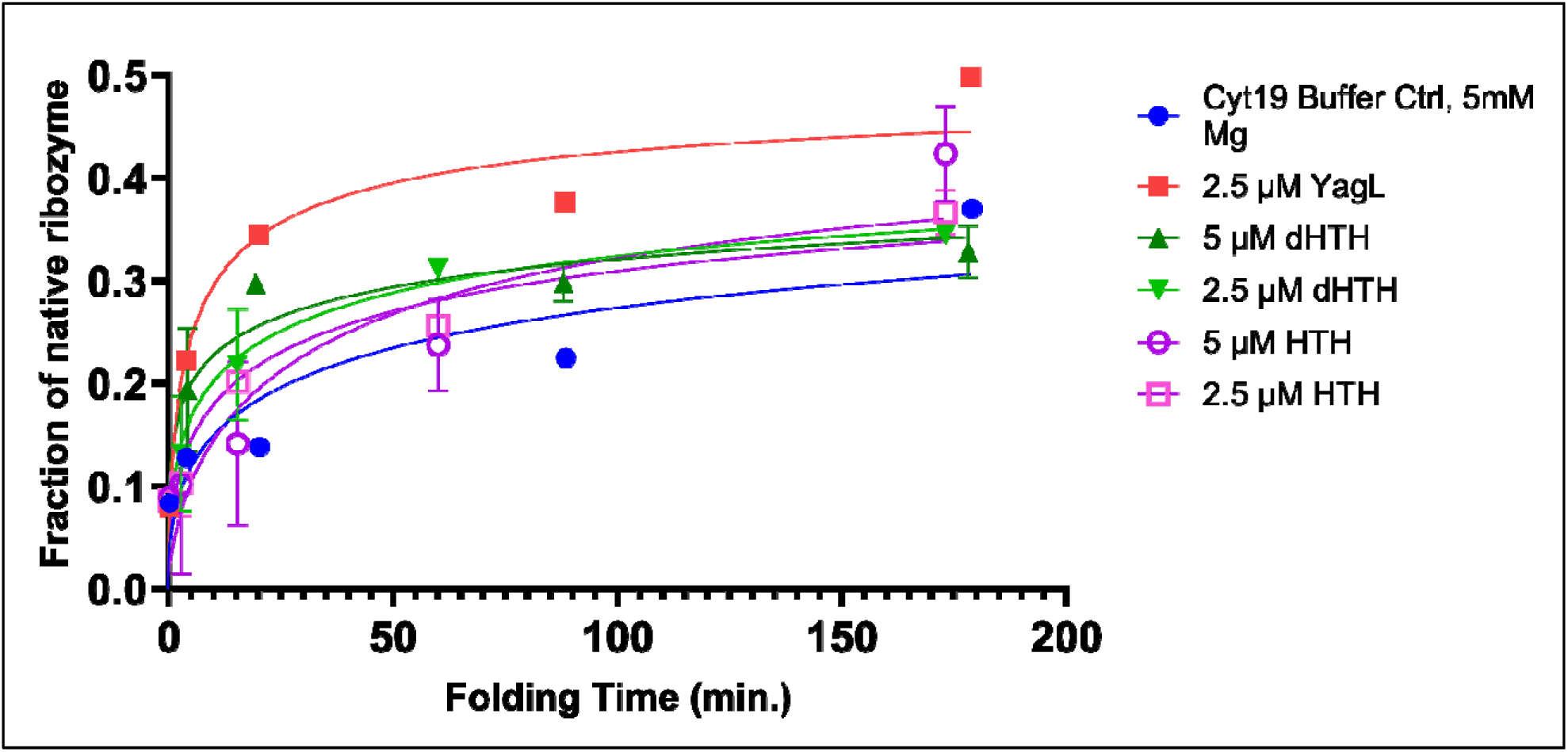
The full YagL protein is required for it to accelerate re-folding of the *Tetrahymena* ribozyme. A two-step catalytic activity assay was performed by pre-incubating the ribozyme into a misfolded state and then adding different concentrations of the “dHTH” or the “HTH” protein truncations. The full length YagL protein was used as a positive control. Reactions were stopped at different times, after which radiolabeled rSA5 was added to perform substrate cleavage reactions. The fraction of cleaved product was quantified and used to generate plots of the fraction of native ribozyme as a function of folding time. Error bars represent the variation between experimental duplicates. Only the full YagL protein was capable of significantly accelerating re-folding of the ribozyme, as evidenced by the rapid increase in fraction of native ribozyme at earlier time points (red) compared to the buffer only control (blue).

## Discussion

The diverse functions of RNAs depend on their folding into precise, native structures (**67**). General RNA chaperones allow folding to occur in biologically relevant time scales by interacting transiently with RNA substrates and resolving non-native intermediate structures (**68, 69**). However, the lack of a shared structural or sequence motif has limited their discovery and characterization. Here, we repurposed a regional accessibility assay (iRS^3^) to identify 31 candidate RNA chaperone proteins (**Table 1**), and we validated *in vivo* effects upon gene deletion for eight of them (*sbcC, pepA, yglW, yagL, argF, yifK, fxsA* and *yjhE)* (**Fig. 3**). We further purified and characterized two of them, YagL and PepA, demonstrating RNA binding activity for both proteins (**Fig. 6 & 7**) and *bona fide* RNA chaperone activity for YagL (**Fig. 5**).

For both the *in vivo* and *in vitro* probing of chaperone activity, we used the well-characterized *Tetrahymena* group I intron ribozyme as an example of a highly structured RNA that is prone to misfolding and benefits from general RNA chaperone activity. Thus, we anticipate that the chaperone action of these proteins will extend to other structured RNAs that require assistance in their folding.

The eight candidate RNA chaperone proteins identified in this study (*sbcC, pepA, yglW, yagL, argF, yifK, fxsA* and *yjhE)* add to the growing list of proteins with demonstrated (or hypothesized) chaperone activity. The large number of proteins with general chaperone activity may reflect that this activity can arise simply from preferential protein binding and thereby stabilization of unfolded or partially unfolded RNA intermediates. Consistent with this idea, RNA chaperone proteins frequently have other functions and act secondarily as chaperones, a role termed ‘moonlighting’ (**72**). Indeed, among the eight chaperone candidates for which we validated ribozyme accessibility increases *in vivo*, we identified several genes encoding predicted and/or known DNA-binding proteins (i.e., *pepA, sbcC, yagL*), and proteins whose functions include DNA-binding have been shown previously to be good candidates for moonlighting as general RNA chaperones (**73, 74**).

It is notable that many known RNA chaperone proteins were not detected in our high-throughput screen. Most generally, lack of detection of a chaperone protein could result from functional redundancy with respect to ribozyme accessibility – i.e., the two proteins facilitate the same structural transitions that give changes in accessibility, and even with one gene deleted, the activity conferred by the other protein is present at a saturating level. Note that such a result would not necessarily indicate that the two proteins are functionally redundant for folding of all RNAs, just the one being probed. It is also possible that a given protein could go undetected under one set of growth conditions while being readily detectable under other growth conditions due to changes in protein level and/or the range or levels of RNA substrates requiring chaperone activity. Additionally, chaperone proteins are unlikely to be detected using our experimental workflow if they make a large contribution to cellular fitness. This is due to sub-culturing and pooling steps preceding sequencing that could cause some mutants to be lost, favoring mutations that are less detrimental to the organism. This might explain why we did not detect the known chaperone StpA in the high-throughput assay, despite finding that an individual deletion of *stpA* resulted in accessibility changes that were similar to those observed upon deletion of *pepA* or *yagL*.

Our biochemical analysis of YagL and PepA revealed properties that strongly support the hypothesis of general RNA chaperone function for these two proteins. Both proteins bind ssRNA with affinities in the low µM range (**Fig. 6 & 7).** ssRNA binding activity is expected to be a near-universal feature of RNA chaperones because the ability of chaperones to accelerate folding transitions requires, by definition, that they bind preferentially and thereby stabilize unfolded intermediates, which are likely to include regions of ssRNA. YagL and PepA also bind dsRNA with similar affinity. This activity may contribute to chaperone activity by trapping helical segments of structured RNA as they transiently unfold from tertiary contacts (**75**). Alternatively, or in addition, dsRNA binding activity may reflect ‘collateral’ effects of the additional functions of these proteins, as PepA binds DNA in Xer site-specific recombination and transcription regulation, and YagL is predicted to function as a resolvase from homology modeling. The relatively modest affinity for RNA appears to be an emerging general property of RNA chaperone proteins, which may reflect the necessity to form a complex that is sufficiently stable to accelerate RNA unfolding but sufficiently transient that it allows for efficient release and subsequent folding of the RNA. Thus, while ATP-dependent chaperones (DEAD-box helicase proteins) use their ATPase activity to cycle between states that have high or low affinity for ssRNA (**14, 76**), ATP-independent chaperones may need to ‘thread the needle’ by having intermediate RNA affinity.

Our biochemical experiments also show directly that YagL possesses robust chaperone activity, as it strongly accelerates refolding of the misfolded *Tetrahymena* ribozyme. This chaperone activity requires the full-length YagL protein, despite the RNA binding and multimerization activities being primarily dependent on the HTH domain **(Fig. 8).** Interestingly, PepA does not possess detectable activity for accelerating this refolding process, yet it does impact accessibility of regions of the ribozyme *in vivo* (those complementary to probes 3, 6, 7 and 8, **Fig. 4A**). Likewise, StpA lacks detectable activity for *Tetrahymena* ribozyme refolding (**Supplementary Fig. S9**) despite affecting accessibility of the ribozyme *in vivo* (**Fig. 1**) and previous demonstrations of chaperone activity for another group I intron and model RNA substrates (**10, 48**). Together, these results highlight that different chaperone proteins have apparently achieved some degree of specialization to the types of RNA folding transitions that they accelerate. This specialization may reflect differences in properties such as RNA binding affinity, specificity, and ability to multimerize; and it probably contributes to our finding that several different chaperone proteins function in *in vivo* folding of a complex, structured RNA like the *Tetrahymena* ribozyme.

Our findings provide initial insights into the chaperone activities of YagL and PepA. In future work, it will be interesting to further investigate their mechanisms of RNA recognition and to elucidate the repertoire of native substrates for these two proteins, as well as for the other candidates that emerged from the high-throughput screen. Additionally, our work illustrates how the iRS^3^ assay could be applied to study *in vivo* folding and chaperone activity in the context of other structured RNAs.

## Additional Information

### Supplementary Material

Supplementary data for this article can be accessed online at XXXXXXX.

### Data Availability Statement

All outputs from Phyre2 simulations and HADDOCK outputs are available in Figshare, at 10.6084/m9.figshare.21899445.

Whole Genome Sequencing data for the strains used in this study is available from the NCBI Sequence Read Archive via BioProject accession number PRJNA923723.

## Acknowledgments

We would like to thank Jessica Podnar and Dennis Wylie (GSAF core of The University of Texas at Austin) for their help in planning the sequencing strategy for the transposon library. We also thank Richard Salinas (Microscopy and Flow Cytometry core of the University of Texas at Austin) for his continuous advice during the flow cytometry experiments. Authors would also like to acknowledge Dr. Maria Person, Michelle Gadush, and the Mass Spectrometry Facility at the University of Texas at Austin for helping in performing Mass Spectrometry Protein Identification experiments. We acknowledge Dr. Jeff Barrick for kindly providing us with the Keio collection deletion strains, Dr. Phanourios Tamamis for providing advice and guidance regarding the molecular docking experiments, and Dr. Mia Mihailovic for insightful discussions and advice on the iRS^3^ experiments. The authors would like to acknowledge BioRender.com for assistance with figure creation. Molecular graphics and analyses performed with UCSF ChimeraX, developed by the Resource for Biocomputing, Visualization, and Informatics at the University of California, San Francisco, with support from National Institutes of Health R01-GM129325 and the Office of Cyber Infrastructure and Computational Biology, National Institute of Allergy and Infectious Diseases.

## Author contributions

Designed Research: A.M.R.N., L.G.M., L.M.C., R.R.; Performed experiments: A.M.R.N., L.G.M., A.A., A.T.M., S.H.J., E.K.; Analyzed data: A.M.R.N., L.G.M.; Wrote paper: A.M.R.N., L.G.M., L.M.C., R.R.

## Funding

This work was supported by the Welch Foundation [F-1756]; NIH [R35 Grant GM131777 to R.R and R01 Grant GM135495 to LMC];] and the Fulbright Program [Fulbright García-Robles to A.M.R.N]. Funding for open access charge: Welch Foundation [F-1756].

## Disclosure statement

No potential conflict of interest was reported by the authors.

## Notes

### Competing Interest Statement

The authors have declared no competing interest.

