## Supplementary Information for "A high-throughput search for intracellular factors that affect RNA folding identifies *E. coli* proteins PepA and YagL as RNA chaperones that promote RNA remodeling"

### Table of Contents

**Supplementary Methods.**

- **Peptidase activity assay.**
- **Bioinformatic analysis of protein structures and amino acid sequences.**
- **DNA ligand selection and molecular docking with the YagL AlphaFold structure.**

**Supplementary Discussion.**

**Supplementary Figures.**

**Supplementary Tables.**

**Appendices.**

**Supplementary References.**

**Supplementary Methods.**

***Peptidase activity assay.***

PepA is a peptidase that catalyzes the release of N-terminal Asp from substrates. To determine the activity of our protein after purification, we performed a catalytic activity assay as described in (**1**) with a few modifications. In this assay, H-Asp-pNA was used as a substrate. 10 uL of $CoCl_2$ (15 mM) were added to 200 µL of purified PepA in storage buffer (Na/K-phosphate buffer; 50 mM; pH 6.0). 200 µL of storage buffer without protein were used as negative control.

After incubating for 10 minutes at 60°C, 12.5 µL of the H-Asp-pNA substrate solution (5 mg/mL $_{DMSO}{}^{-1}$) were added to the reaction mixture. The reaction was terminated after 5 minutes by adding 50 µL of acetic acid (50% (v/v) to the sample. Reactions were centrifuged (8000 x g, 5 minutes, 4°C) and 240 µL of the supernatant were transferred into a 96-well plate. Absorption was measured at 405 nm. Activity was determined using a standard curve with known concentrations of p-NA **(Supplementary Fig. S3).** The amount of pNA released was used to calculate the activity of PepA in katals (kat), where kat is defined as the release of 1 mol p-nitroanilin per s (**1**).

***Bioinformatic analysis of protein structures and amino acid sequences.***

To further investigate the structural composition of the YagL and PepA proteins, structural predictions were generated for their amino acid sequences using the Phyre2 suite of tools. Phyre2 (https://www.sbg.bio.ic.ac.uk/phyre2/) is a fold recognition and structure prediction server that generates a Hidden Markov Model to compare a protein query against published protein structures (**2**). Sequences were submitted to the webserver under the intensive mode and the generated structures for both the entire protein and relevant subdomains were visualized in UCSF ChimeraX (**3**).

In addition to a homology modeling approach with Phyre2, the newly developed AlphaFold2 was also used to generate a predicted structure for YagL, since the structure for this protein has yet to be resolved (**4**). AlphaFold uses a deep learning algorithm to develop a structural prediction, based on a multiple sequence alignment (MSA) representation, that rivals the accuracy of experimental methods. The structure from the AlphaFold Protein Structure Database for YagL was used for further conservation analysis and computational docking simulations (**5**).

The ConSurf web server was used to visualize the relative conservation of different amino acid residues within the predicted YagL structure and, independently, the published PepA crystal structure (PDB:1GYT) (**6-9**). Briefly, the AlphaFold structure for YagL and the crystallized PepA were provided as input to the server. The protein sequences were then extracted from the PDB and BLASTed against proteins in the UNIREF-90 database. After further filtering this list of aligned proteins, conservation analysis was performed to yield per-residue grading for conservation on a scale from 1 to 9, with 9 showing the highest relative conservation at that position (**2**). These gradings were color coded and mapped onto the protein structures. The ConSurf structure outputs were then visualized with UCSF ChimeraX (**3**). ChimeraX was also used to visualize the qualitative electronegativity landscape of each structure with electrostatics calculated using the Adaptive Poisson-Boltzmann Solver (**10**).

***DNA ligand selection and molecular docking with the YagL AlphaFold structure.***

To further investigate which regions in YagL might contribute to nucleic-acid binding and possibly RNA chaperone activity, six DNA ligands were chosen from the list of homologous protein structures identified from the Phyre2 modeling output. These DNA structures, along with the YagL structure generated with AlphaFold, served as input for molecular docking simulations using the HADDOCK webserver (**11-13**). HADDOCK uses a data-driven approach to predict accurate docking poses, and HADDOCK2.2 was used for its support for mixed (i.e., protein-DNA) docking (**14, 15**). The first 46 residues from the N-terminus of the YagL structure were defined as passive residues in the docking setup due to their lower confidence structure in both AlphaFold2 and Phyre2 outputs. Standard docking settings for simulations including nucleic acids were used. Ambiguous interaction restraints (AIRs) were randomly excluded at a rate of 50%. 1000 structures were generated for rigid body docking, 200 for semi-flexible refinement, and 200 for the final refinement. The RMSD clustering method was used for these simulations, with a cutoff of 7.5 angstroms. Clustered structures were ranked by their HADDOCK score, which is calculated as a combination of energetic terms (**15**). Representative structures from the best-scoring clusters were subjected to further refinement using the HADDOCK web server and visualized using ChimeraX (**3**).

**Supplemental Discussion.**

***Bioinformatic analysis and molecular docking simulations.***

To identify possible molecular features responsible for RNA chaperone activity, the predicted structure of YagL was examined using bioinformatic and computational methods. The predicted structure for YagL shows similarity to resolvases, which represent a specific type of recombinase that interacts with DNA through covalent interactions using the N-terminus domain of the protein and a series of contacts with the minor and major grooves of DNA mediated by the HTH at the C-terminus domain (**16**). Specifically, a resolvase of a similar topology has been proposed to resolve supercoiled, circular DNA into two catenated molecules via double-strand DNA cleavage, strand exchange, and re-ligation (**17**). To identify key molecular features that might be implicated in the affinity of YagL for DNA, and possibly its RNA substrate by extension, we performed molecular docking simulations using the HADDOCK webserver (**11-13**). To serve as input, we chose six DNA chains from homologous resolvase crystal structures identified in the Phyre2 output for YagL (**Supplementary Table S3**) along with the AlphaFold predicted structure for YagL. Following modeling and refinement, the protein-DNA interfaces for all 6 models were examined using a 7.5 angstrom distance cutoff (**Supplemental Fig. S8**). Nucleic-acid interfacing residues within the YagL HTH domain are defined as residues that appear within the distance cutoff for at least 5 of the 6 docked complex structures. Residues such as histidine, lysine, arginine, and tryptophan are identified as nucleic acid interfacing residues from the docked structures. These positively charged, polar, or aromatic residues can interact with both the phosphate backbone of DNA and RNA but also with the nucleobases, as can be seen with SRSF1 or RNase T (**18-21**). These residues, in agreement with the mode of binding for known resolvases, are mostly found at the N-terminus domain and within the HTH at the C-terminus domain of YagL and represent interesting candidates for further *in vitro* and *in vivo* studies.

#### Supplementary Figures.

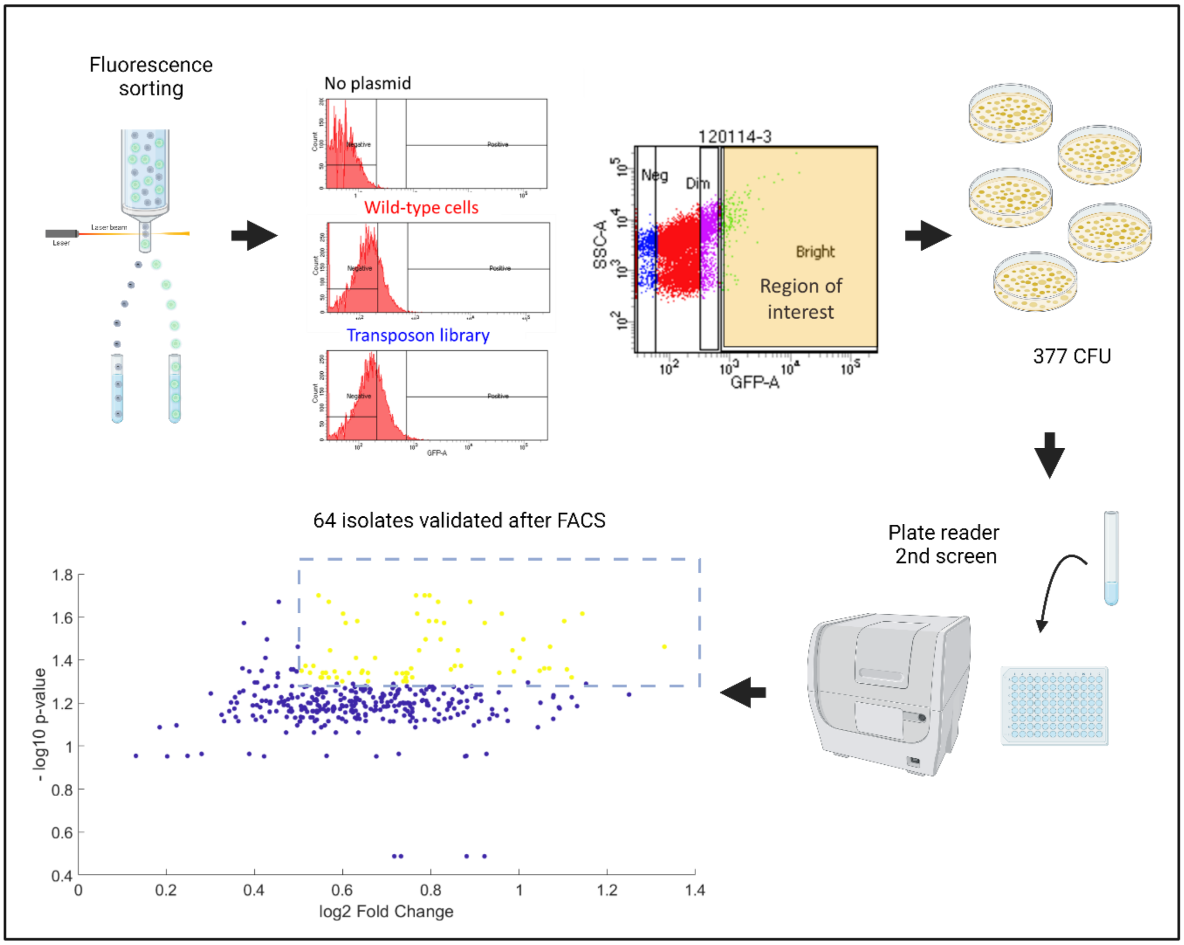


**Supplementary Figure S1. Schematic overview of the fluorescence screening and validation process.** Library collections composed of transposed *E. coli* cells expressing the iRS$^3$ reporter were screened and sorted based on their fluorescence intensity using a FACSAria IIIu cell sorter (Becton Dickinson). Isolates from the region of interest corresponded to 1.1-1.7 % of the total population. After the initial screen and recovery, sorted colonies were screened again in a 96-well plate using a Cytation3 plate reader. Sixty-four isolates were selected for further characterization, using a log2 fold-change cutoff of > 0.5 and a p-value < 0.05. Figure created with BioRender.com.

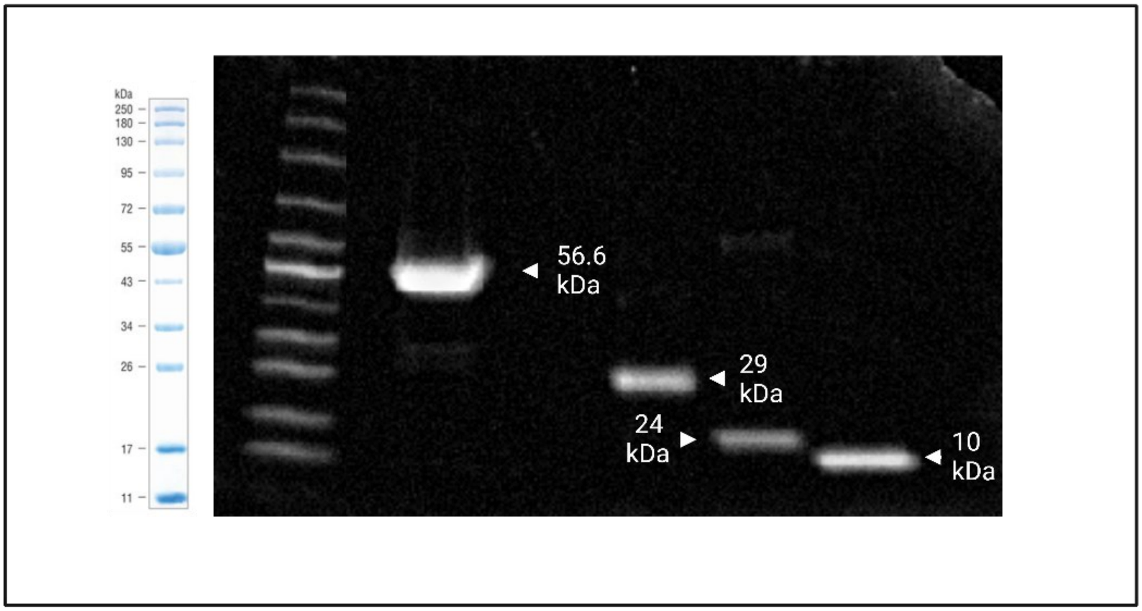


**Supplementary Figure S2. His-tag purification of chaperone proteins.** Proteins were visualized using Coomassie blue staining. Purified PepA was observed as a single band at ~56.6 kDa. YagL was observed as a single band at ~29 kDa. YagL-dHTH was observed as a single band at ~ 24 kDa. YagL-HTH was observed as a single band at ~10 kDa. Protein purity was further validated via LC-MS/MS. No proteins were detected within 10-fold of the purified protein for each sample.

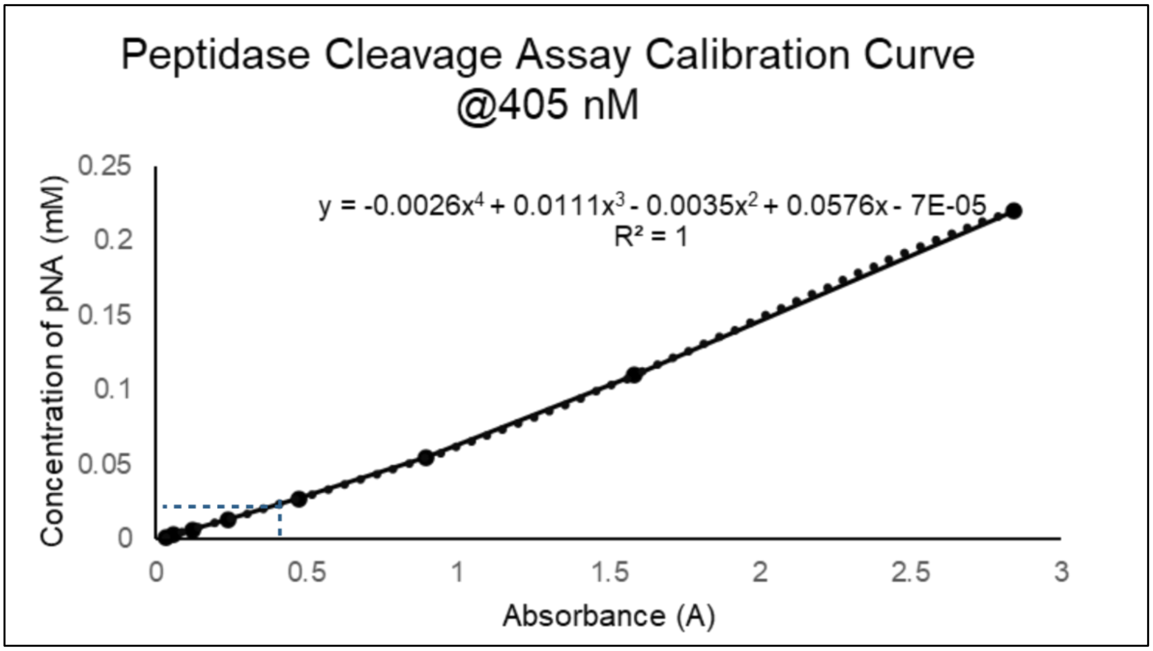

**Supplementary Figure S3. Functionality of PepA after purification was evaluated using a peptidase cleavage assay.** A standard curve was created using known concentrations of pNA and plotting their absorbance at 405 nM. H-Asp-pNA was used as a substrate and the concentration of cleaved product, pNA, after a 10-minute reaction was determined by comparing the absorbance against the standard curve. The specific activity of our purified PepA was determined to be ~73.1 nKat/mg of protein.

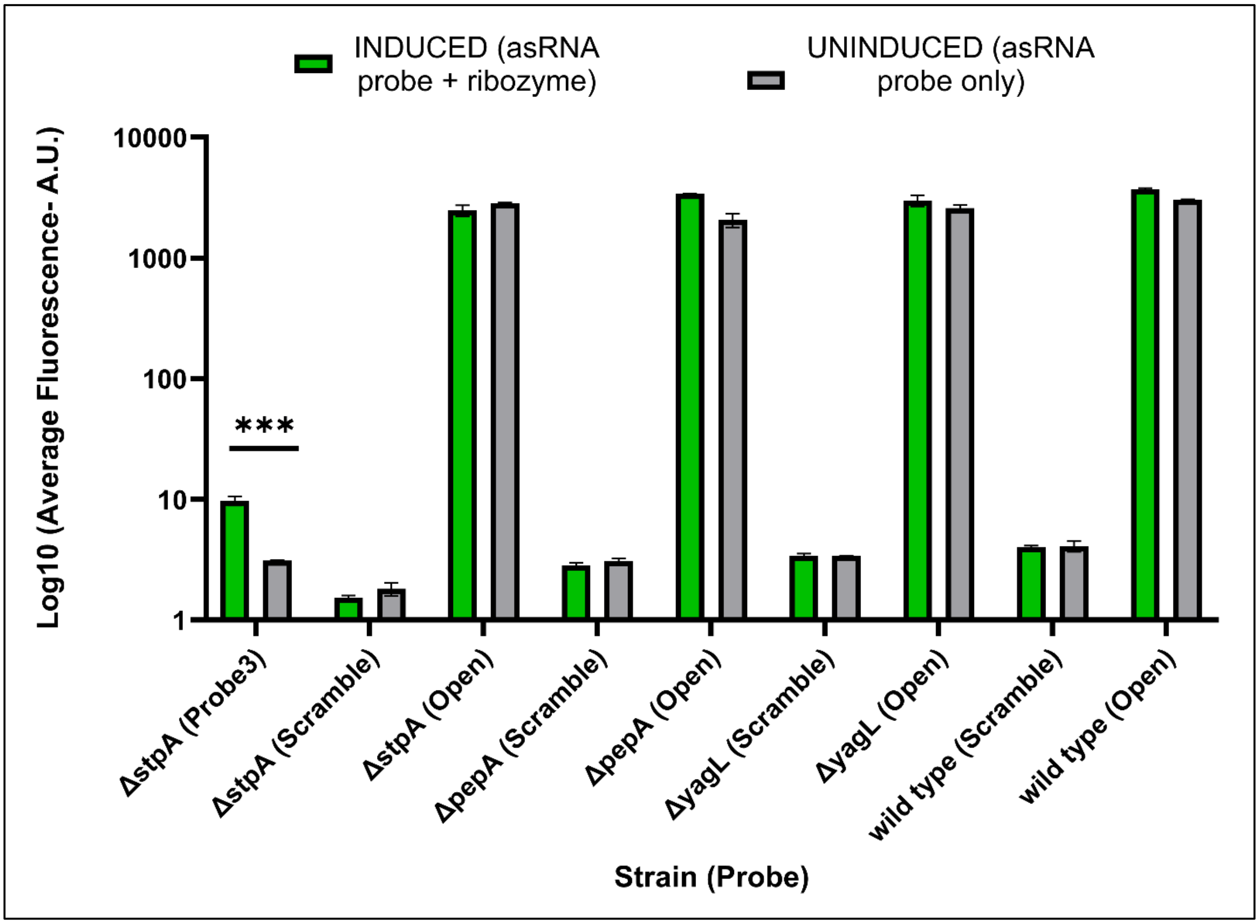


**Supplementary Figure S4. Fluorescence measurements for low throughput iRS$^3$ controls**. iRS$^3$ fluorescence measurements corresponding to Probe 3 and previously published (**22**) control probes "Open" and "Scramble" overexpressed in *E. coli*. The "Open" control represents the high-end of fluorescence since point mutations prevent sequestration of the probe ribosome binding site (RBS) and thus translation is always active. The "Scramble" control contains a random asRNA sequence that is not complementary to the *E. coli* genome and thus should always remain in a closed/no translation conformation, representing the low-end of fluorescence. For each probe, fluorescence average fluorescence was measured upon induction of the ribozyme (INDUCED condition) and with only the asRNA probe being expressed (target-uninduced or UNINDUCED condition). An "Open" and a "Scramble" control were included in every iRS$^3$ experiment as positive and negative fluorescence controls respectively.

**A**

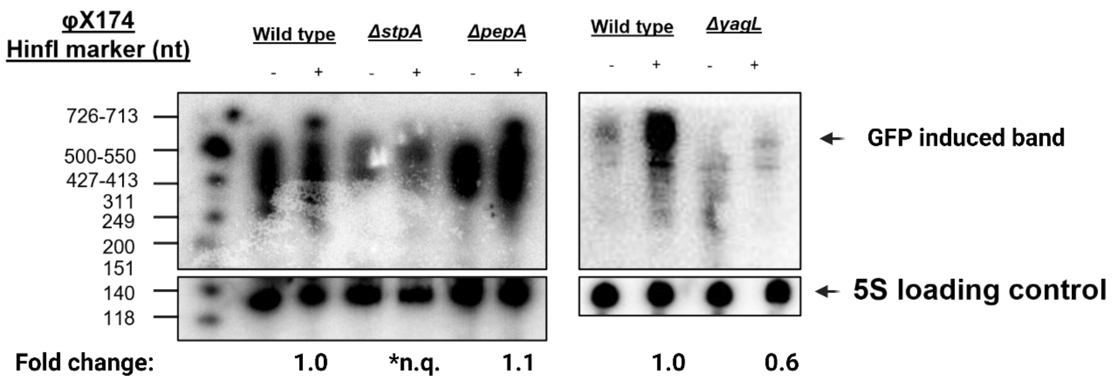


**B**

$$Fold\ change = \frac{Induced\ GFP\ band}{(Induced\ GFP\ band)_{WT}}\ x\ \frac{5S\ loading\ control_{WT}}{5S\ loading\ control}$$

**Supplementary Figure S5. Northern blotting analysis of the iRS$^3$ system.** Total RNA extracted from cultures from the fluorescence assay performed on the *ΔpepA, ΔyagL,* and wildtype *E. coli* K-12 BW25113. RNA extracted from the *ΔstpA* strain was blotted for comparison. The iRS$^3$ probe was blotted using a GFP universal probe (*5'- GCCCATTAACATCACC- 3'*). The 5s ribosomal RNA was blotted as a loading control using probe GII *(5'- TGCATGCCTGATAACTTT -3')*. Quantification (imageJ) was performed, and each band was normalized to the 5S band on the same lane. (B) Relative intensities were calculated and used to determine fold-changes so that the wild type (WT) band equals 1. * Fold change was not calculated for the induced GFP signal of the *ΔstpA* strain on the depicted membrane due to high brightness noise during imaging.

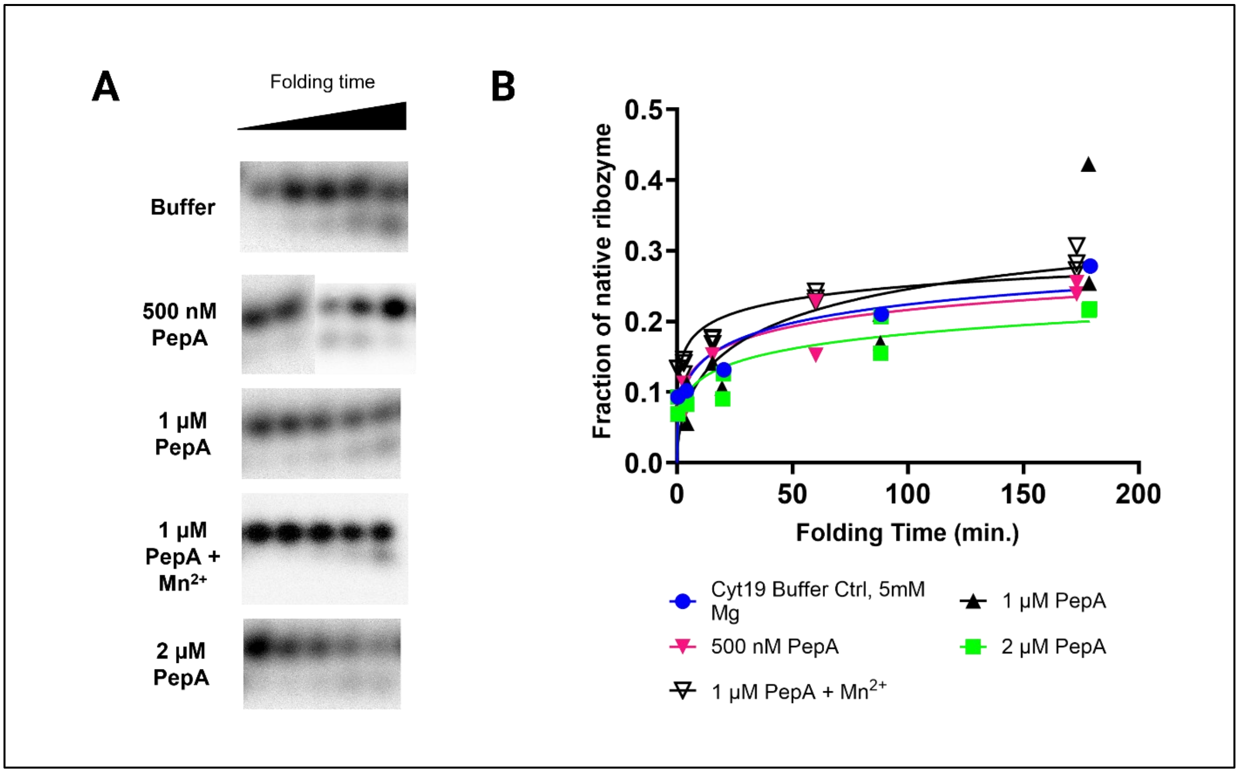


**Supplementary Figure S6. Native ribozyme folding in the presence of PepA** (A) The *Tetrahymena* ribozyme was pre-incubated into a misfolded state and then added into folding reactions without PepA (α), with 500 nM PepA (β), 1 µM PepA (γ), 1 µM PepA + 0.5 mM $Mn^{2+}$ (δ), or 2 µM PepA (ε). Reactions were stopped at different times, after which radiolabeled rSA5 was added to perform substrate cleavage reactions. After quenching, reactions were stopped and loaded into a 20% denaturing PAGE gel to quantify cleavage product formation. Representative gel images of these reactions are shown in this panel. Reactions were performed in independent duplicates (raw and processed images for all experimental runs can be found in **Supplementary Information- Appendix 3).** Due to the number of conditions tested, the reactions with 500 nM PepA were splitted and run simultaneously in two different gels inside the same tank and at the same conditions. The images collected from these two gels are shown side by side. (B) Rate constants for native state formation were 0.0016 min-1 (α), 0.0015 min-1 (β), 0.0016 min-1 (γ), 0.0022 min-1 (δ), and 0.0015 min-1 (ε). No significant differences were observed in the observed rate constants by the addition of PepA under the tested conditions.

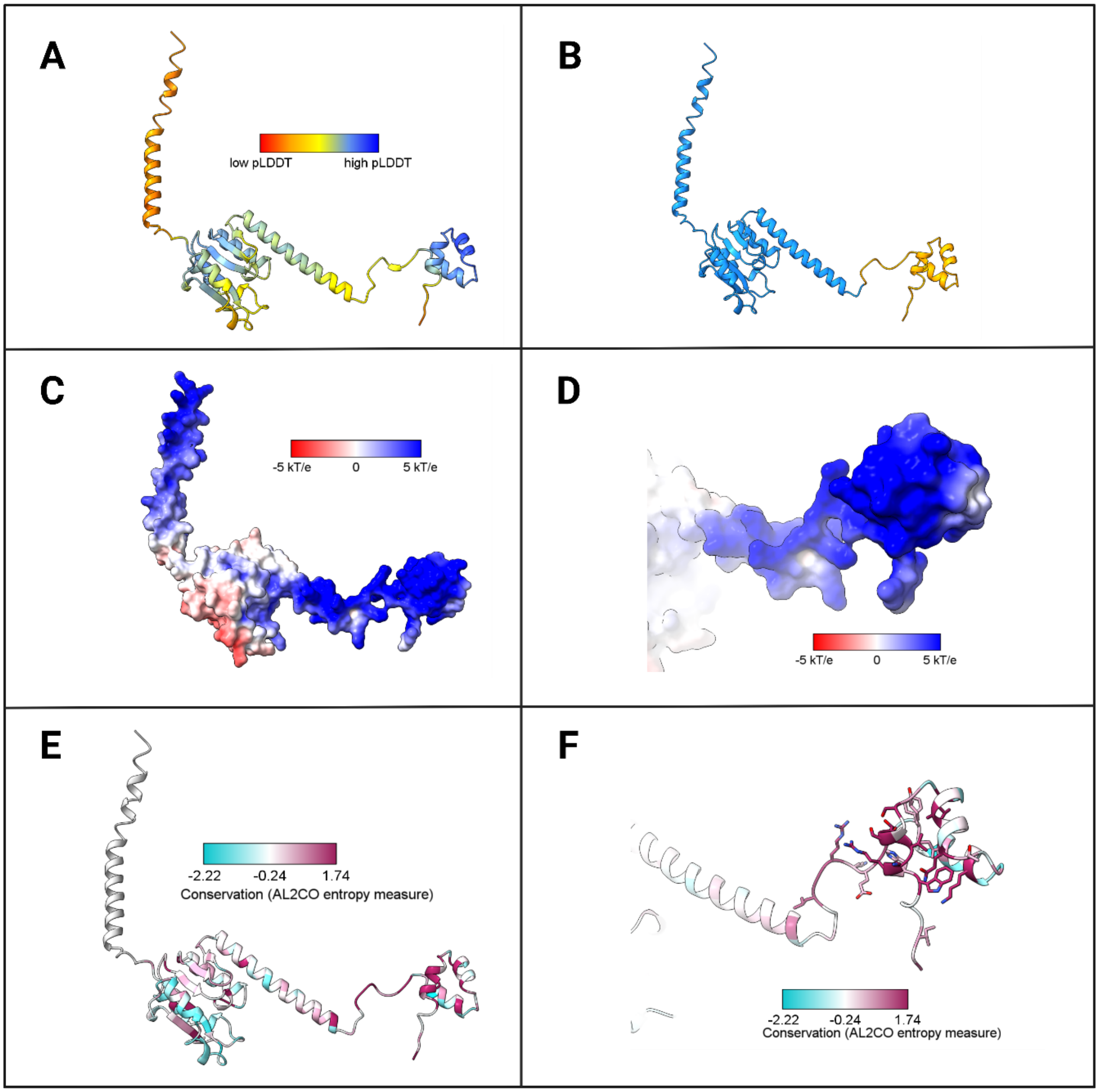


**Supplementary Figure S7. The structural model of YagL contains a C-terminal helix-turn-helix with conserved, positively charged residues.** Molecular visualizations generated with ChimeraX (**3**). (A) The structural model for YagL generated with Alphafold2 (**4**). Residues colored by model confidence using the predicted local distance difference test (pLDDT). (B) The YagL model shows a structured N-terminal domain (blue) and a C-terminal helix-turn-helix domain (orange) predicted with high confidence. (C-D) The relative electrostatics of the YagL model were calculated using the APBS webserver (**10**) showing a region of positively charged residues in the helix-turn-helix region. (E-F) Conservation analysis of the structural model via the CONSURF web server (**6-9**) shows high conservation (high AL2CO entropy measure) of the residues in the helix-turn-helix domain.

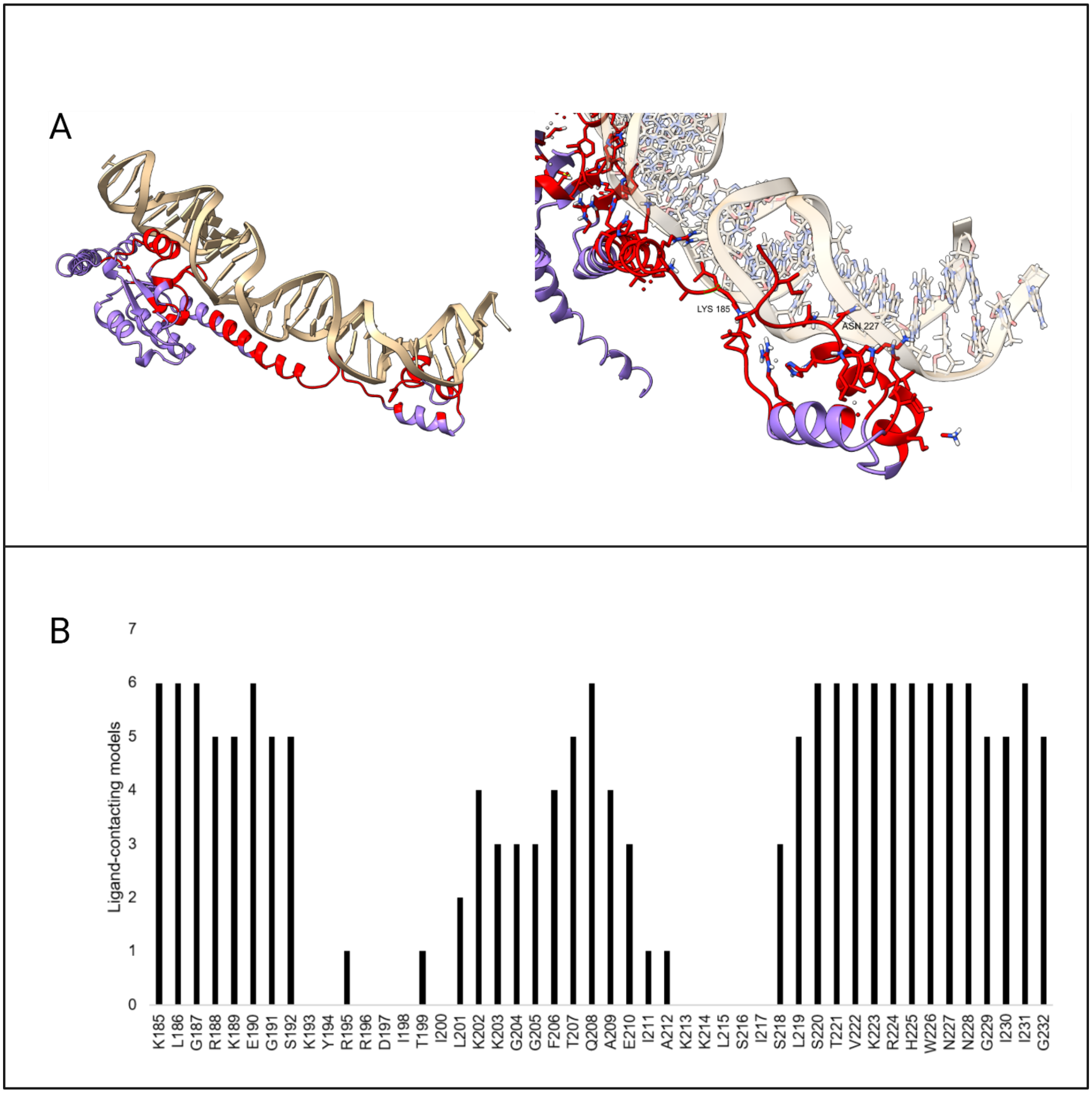


**Supplementary Figure S8. Protein residues in the helix-turn-helix domain commonly interact with DNA across docking simulations.** (A) A representative docking pose is shown (YagL: DNA ligand 01) with residues within 7.5 angstroms of the DNA chain highlighted in red. (B) Frequency of helix-turn-helix residues within the distance cutoff of the DNA chain. Residues predicted to be nucleic acid interfacing are those within the distance cutoff for 5 of the 6 docked structures.

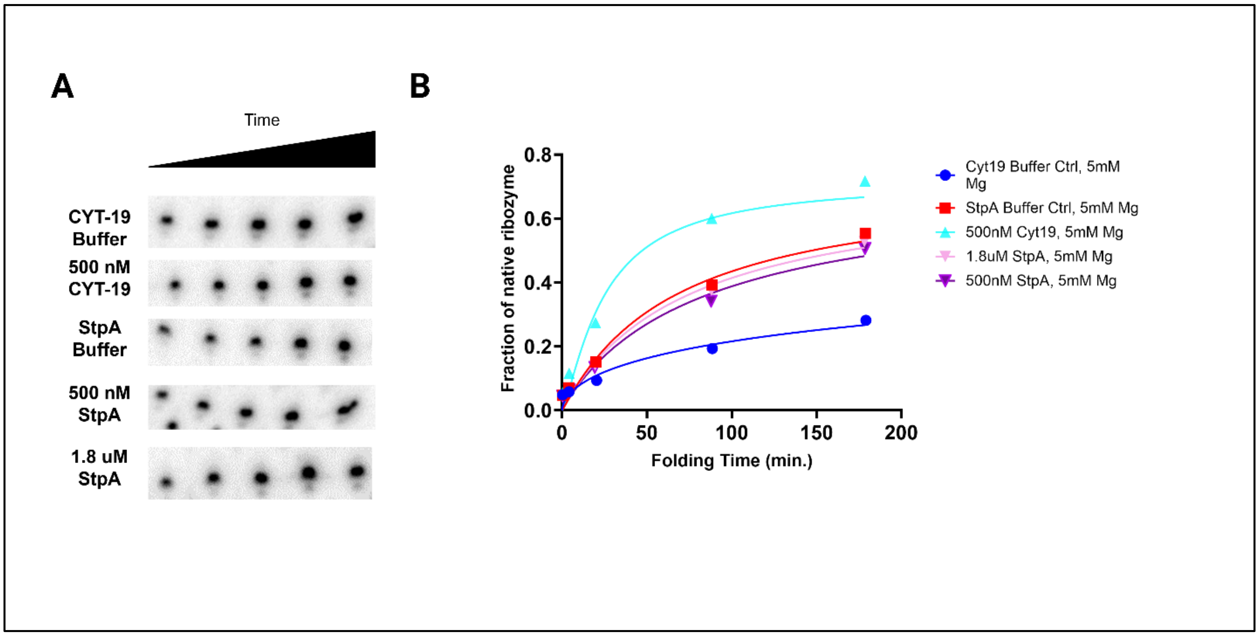


**Supplementary Figure S9. Catalytic activity assays in the presence of CYT-19 and StpA.** (A) The *Tetrahymena* gI intron ribozyme was pre-incubated into a misfolded state and then added into folding reactions without chaperone protein (α), with 500 nM CYT-19 (β), StpA storage buffer (γ), 500 nM StpA, or 1.8 µM StpA (ε). ATP was added to the reactions with CYT-19. Reactions were stopped at different times, after which radiolabeled rSA5 was added to perform substrate cleavage reactions. After quenching, reactions were stopped and loaded into a 20% denaturing PAGE gel to quantify cleavage product formation. (B) Rate constants for native state formation were 0.0024 $min^{-1}$ (α), 0.0187 $min^{-1}$ (β), 0.0083 $min^{-1}$ (γ), 0.0076 $min^{-1}$ (δ), and 0.0068 $min^{-1}$ (ε). No significant differences were observed in these rate constants by the addition of StpA in the tested conditions.

**Supplementary Tables.**

| Plasmid | Strain | Description/Cloning method | Reference |
|---|---|---|---|
| none | *E. coli* MG1655 | | Blattner, F.R. *et al.* (1997) |
| none | *E. coli* BW25113 parent strain | | Baba, T. *et al.* (2006) |
| none | *E. coli* BW25113 *ΔsbcC::kan* | cured strain; kanamycin cassette removed using FLP recombination | Baba, T. *et al.* (2006) |
| none | *E. coli* BW25113 *ΔpepA::kan* | cured strain; kanamycin cassette removed using FLP recombination | Baba, T. *et al.* (2006) |
| none | *E. coli* BW25113 *ΔygiW::kan* | cured strain; kanamycin cassette removed using FLP recombination | Baba, T. *et al.* (2006) |
| none | *E. coli* BW25113 *ΔcyaA::kan* | cured strain; kanamycin cassette removed using FLP recombination | Baba, T. *et al.* (2006) |
| none | *E. coli* BW25113 *ΔyjjL::kan* | cured strain; kanamycin cassette removed using FLP recombination | Baba, T. *et al.* (2006) |
| none | *E. coli* BW25113 *ΔilvL::kan* | cured strain; kanamycin cassette removed using FLP recombination | Baba, T. *et al.* (2006) |
| none | *E. coli* BW25113 *ΔyifK::kan* | cured strain; kanamycin cassette removed using FLP recombination | Baba, T. *et al.* (2006) |
| none | *E. coli* BW25113 *ΔyagL::kan* | cured strain; kanamycin cassette removed using FLP recombination | Baba, T. *et al.* (2006) |
| none | *E. coli* BW25113 *ΔphoB::kan* | cured strain; kanamycin cassette removed using FLP recombination | Baba, T. *et al.* (2006) |
| none | *E. coli* BW25113 *ΔybcL::kan* | cured strain; kanamycin cassette removed using FLP recombination | Baba, T. *et al.* (2006) |
| none | *E. coli* BW25113 *ΔyghQ::kan* | cured strain; kanamycin cassette removed using FLP recombination | Baba, T. *et al.* (2006) |
| none | *E. coli* BW25113 *ΔyhgF::kan* | cured strain; kanamycin cassette removed using FLP recombination | Baba, T. *et al.* (2006) |
| none | *E. coli* BW25113 *ΔyjeM::kan* | cured strain; kanamycin cassette removed using FLP recombination | Baba, T. *et al.* (2006) |
| none | *E. coli* BW25113 *ΔilvC::kan* | cured strain; kanamycin cassette removed using FLP recombination | Baba, T. *et al.* (2006) |

| | | | |
|---|---|---|---|
| none | *E. coli* BW25113 *ΔargF::kan* | cured strain; kanamycin cassette removed using FLP recombination | Baba, T. *et al.* (2006) |
| none | *E. coli* BW25113 *ΔglgX::kan* | cured strain; kanamycin cassette removed using FLP recombination | Baba, T. *et al.* (2006) |
| none | *E. coli* BW25113 *ΔyjcC::kan* | cured strain; kanamycin cassette removed using FLP recombination | Baba, T. *et al.* (2006) |
| none | *E. coli* BW25113 *ΔpanF::kan* | cured strain; kanamycin cassette removed using FLP recombination | Baba, T. *et al.* (2006) |
| none | *E. coli* BW25113 *ΔbcsC::kan* | cured strain; kanamycin cassette removed using FLP recombination | Baba, T. *et al.* (2006) |
| none | *E. coli* BW25113 *ΔfxsA::kan* | cured strain; kanamycin cassette removed using FLP recombination | Baba, T. *et al.* (2006) |
| none | *E. coli* BW25113 *ΔyjhE::kan* | cured strain; kanamycin cassette removed using FLP recombination | Baba, T. *et al.* (2006) |
| pACYC184 | | | Rose, R.E. (1988) |
| pMOD2 | | | Epicentre Biotechnology |
| pCML375 | | pMOD2 derivative; TetR cloned from pACYC184 vector using HindIII and EcoRI cut sites, PCR amplified | present study |
| pCYFP | | | Rojano-Nisimura, A.M. *et al.* (2020) |
| pCML2868 | pET21-stpA | pCYFP derivative; stpA amplified from the genome with homology arms and cloned into pET21 by Gibson Assembly | present study |
| pCML2870 | pET21-pepA | pCYFP derivative; pepA amplified from the genome with homology arms and cloned into pET21 by Gibson Assembly | present study |
| pCML3724 | pET21-yagL | pCYFP derivative; YagL amplified from the genome with homology arms and cloned into pET21 by Gibson Assembly | present study |
| pCML2531 | p-OiRS3GG-p1 | wild-type group I intron (193) and probe 218 (probe 1) inserted between EcoRI site and start of CB region | Sowa, S.W. *et al.* (2014) |
| pCML2532 | p-OiRS3GG-p2 | wild-type group I intron (193) and probe 785 (probe 2) inserted between EcoRI site and start of CB region | Sowa, S.W. *et al.* (2014) |
| pCML2533 | p-OiRS3GG-p3 | wild-type group I intron (193) and probe 250 (probe 3) inserted between EcoRI site and start of CB region | Sowa, S.W. *et al.* (2014) |
| pCML2534 | p-OiRS3GG-p4 | wild-type group I intron (193) and probe 786 (probe 4) inserted between EcoRI site and start of CB region | Sowa, S.W. *et al.* (2014) |
| pCML2535 | p-OiRS3GG-p5 | wild-type group I intron (193) and probe 787 (probe 5) inserted between EcoRI site and start of CB region | Sowa, S.W. *et al.* (2014) |
| pCML2536 | p-OiRS3GG-p6 | wild-type group I intron (193) and probe 221 (probe 6) inserted between EcoRI site and start of CB region | Sowa, S.W. *et al.* (2014) |
| pCML2537 | p-OiRS3GG-p7 | wild-type group I intron (193) and probe 788 (probe 7) inserted between EcoRI site and start of CB region | Sowa, S.W. *et al.* (2014) |
| pCML2538 | p-OiRS3GG-p8 | wild-type group I intron (193) and probe 219 (probe 8) inserted between EcoRI site and start of CB region | Sowa, S.W. *et al.* (2014) |

| | | | |
|---|---|---|---|
| pCML698 | p-OiRS3GG-p9 | wild-type group I intron (193) and probe 223 (probe 9) inserted between EcoRI site and start of CB region | Sowa, S.W. *et al.* (2014) |
| pCML2539 | p-OiRS3GG-p10 | wild-type group I intron (193) and probe 789 (probe 10) inserted between EcoRI site and start of CB region | Sowa, S.W. *et al.* (2014) |
| pCML1927 | p-O-iRS3GG-scramble | 15 bp scramble asRNA (CAGCGACAATATCGT) | Leistra, A.N., Mihailovic, M.K. *et al.* (2018) |
| pCML1928 | p-O-iRS3GG-open | free RBS: scramble with mutated CB (GCATAAATTAGGGAGTCAA) | Leistra, A.N., Mihailovic, M.K. *et al.* (2018) |
| pCML3725 | pET21-YagL-ΔHTH | coding region from AA 1-186 amplified from pET21-yagL; inserted into pET21 backbone between StyI site and XhoI site by Gibson Assembly<br>primers: backbone_fwd (GACATGTTAAGAAAAAATTACTCGAGCACCACCACCAC), backbone_rev (GGATCCTCCCTTGTCGTC)<br>insert_fwd (ATGACGACAAGGGAGGATCCAGAGGTAAAATACTACTTTACC), insert_rev (TAATTTTTTCTTAACATGTCGATATC) | present study |
| pCML3739 | pET21-YagL-HTH | coding region from AA 187-232 amplified from pET21-yagL; inserted into pET21 backbone between StyI site and XhoI site by Gibson Assembly<br>primers: backbone_fwd (GGAATAACGGAATCATCGGTCTCGAGCACCACCACCAC), backbone_rev (GGATCCTCCCTTGTCGTC)<br>insert_fwd (ATGACGACAAGGGAGGATCCGGACGAAAGGAGGGAAGC), insert_rev (ACCGATGATTCCGTTATTCC) | present study |

**Supplementary Table S1. List of strains, plasmids and cloning techniques used in this study.**

| | | |
|---|---|---|
| gI intron ribozyme insert | 393 nt | GGAGGGAAAAGTTATCAGGCATGCACCTGGTAGCTAGTCTTTAAACCAATAGATTGCATCGGTTTAAAAGGCAAGACCGTCAAATTGCGGGAAAGGGGTCAACAGCCGTTCAGTACCAAGTCTCAGGGGAAACTTTGAGATGGCCTTGCAAAGGGTATGGTAATAAGCTGACGGACATGGTCCTAACCACGCAGCCAAGTCCTAAGTCAACAGATCTTCTGTTGATATGGATGCAGTTCACAGACTAAATGTCGGTCGGGGAAGATGTATTCTTCTCATAAGATATAGTCGGACCTCTCCTTAATGGGAGCTAGCGGATGAAGTGATGCAACACTGGAGCCGCTGGGAACTAATTTGTATGCGAAAGTATATTGATTAGTTTTGGAGTACTCG |

**Supplementary Table S2. Sequence of the *Tetrahymena* gI intron ribozyme insert present in pCML375 (Addgene number: (Plasmid #98589)).**

| Identity for this study | Structure description | DNA chain(s) used for docking | Reference |
|---|---|---|---|
| Ligand 1 | Crystal Structure of a serine recombinase- DNA regulatory complex | 2R0Q:B | 2R0Q (PMID: 18439894) |
| Ligand 2 | Crystal Structure of a site-specific recombinase, Gamma-delta resolvase complexed with a 34 bp cleavage site | 1GDT:A, 1GDT:B, 1GDT:D, 1GDT:C | 1GDT (PMID: 7628011) |
| Ligand 3 | DeoR protein bound to DNA operator | 7BHY:A, 7BHY:B | 7BHY (PMID: 34726169) |
| Ligand 4 | Pax5Ets Binding Site on the mb-1 promoter | 1K78:C, 1K78:D | 1K78 (PMID: 11779502) |
| Ligand 5 | Human Pax6-protein bound to its DNA-binding domain | 6PAX:A, 6PAX:B | 6PAX (PMID: 10346815) |
| Ligand 6 | Hin recombinase bound to half of a DNA recombination site | 1HCR:A, 1HCR:B | 1HCR (PMID: 8278807) |

**Supplementary Table S3. DNA ligand structures for computational docking with YagL.** Ligands were chosen from their parent structures due to homology identified from Phyre2 output. The structures of the DNA chains were isolated from their parent models and used as input for molecular docking simulations with the YagL structural model.

| | | |
|---|---|---|
| YagL full sequence | 759 bp | ATGGCCGACTACAAGGACGATGACGACAAGGGAGGATCCAGAGGTAAAATA<br>CTACTTTACCAGTTAAAGTATCGCTGGCAATCACTCTCTATTTTTGGTTGTTTT<br>TTATGCAAAATGACATTATTCAGGTATCAAAAAATAATTTATGACACCGGAG<br>TGCATCAGATGCGTTCATTTTTTTACACCATATGCAGTAGCGAACAGCAAGA<br>AAGCATTACAGATCATCACTCACTTGCTGAGATATGCCAAAAATTTAATATT<br>CTACCTGAACATGTTGTGATCGAGCAGGTAGATATTAAAGAGGTTGTTTCAG<br>AGCAACGTTTACTCAGACAATTGATTCATCATGAAATGAACCGGCAGGACAC<br>GCTTGTCATTCCAGATCTAAGCTGTCTTGGCAGAACTGTGGAGGACTTACAG<br>AATATTTTATTTTTTTGCCTTCAAAAAGAAATGTTTATCTACAGCTACCATCC<br>AGCTTCAAGGATAGAGCCTTCTGCTGAAAGTTGCTTGTCTTTTTTGATTGCTC<br>GTCAGGATACGATTGACATTCACAATCTGAAATCAACTAAAAGCCGATATCG<br>ACATGTTAAGAAAAAAATTAGGACGAAAGGAGGGAAGCAAATACCGACGTGA<br>TATCACAATACTTAAGAAAGGTGGATTTACTCAGGCTGAGATTGCAAAGAAA<br>TTGAGTATCAGTTTATCGACTGTGAAGCGGCACTGGAATAACGGAATCATCG<br>GTctcgagcaccaccaccaccaccactga |

| | | |
|---|---|---|
| YagL-dHTH | 621 bp | ATGGCCGACTACAAGGACGATGACGACAAGGGAGGATCCAGAGGTAAAATA<br>CTACTTTACCAGTTAAAGTATCGCTGGCAATCACTCTCTATTTTTGGTTGTTTT<br>TTATGCAAAATGACATTATTCAGGTATCAAAAAATAATTTATGACACCGGAG<br>TGCATCAGATGCGTTCATTTTTTTACACCATATGCAGTAGCGAACAGCAAGA<br>AAGCATTACAGATCATCACTCACTTGCTGAGATATGCCAAAAATTTAATATT<br>CTACCTGAACATGTTGTGATCGAGCAGGTAGATATTAAAGAGGTTGTTTCAG<br>AGCAACGTTTACTCAGACAATTGATTCATCATGAAATGAACCGGCAGGACAC<br>GCTTGTCATTCCAGATCTAAGCTGTCTTGGCAGAACTGTGGAGGACTTACAG<br>AATATTTTATTTTTTTGCCTTCAAAAAGAAATGTTTATCTACAGCTACCATCC<br>AGCTTCAAGGATAGAGCCTTCTGCTGAAAGTTGCTTGTCTTTTTTGATTGCTC<br>GTCAGGATACGATTGACATTCACAATCTGAAATCAACTAAAAGCCGATATCG<br>ACATGTTAAGAAAAAATTActcgagcaccaccaccaccaccactga |
| HTH | 204 bp | ATGGCCGACTACAAGGACGATGACGACAAGGGAGGATCCGGACGAAAGGA<br>GGGAAGCAAATACCGACGTGATATCACAATACTTAAGAAAGGTGGATTTACT<br>CAGGCTGAGATTGCAAAGAAATTGAGTATCAGTTTATCGACTGTGAAGCGGC<br>ACTGGAATAACGGAATCATCGGTctcgagcaccaccaccaccaccactga |

**Supplementary Table S4. YagL protein truncations sequences.**

**Appendix 1. Phyre2 outputs for YagL.**

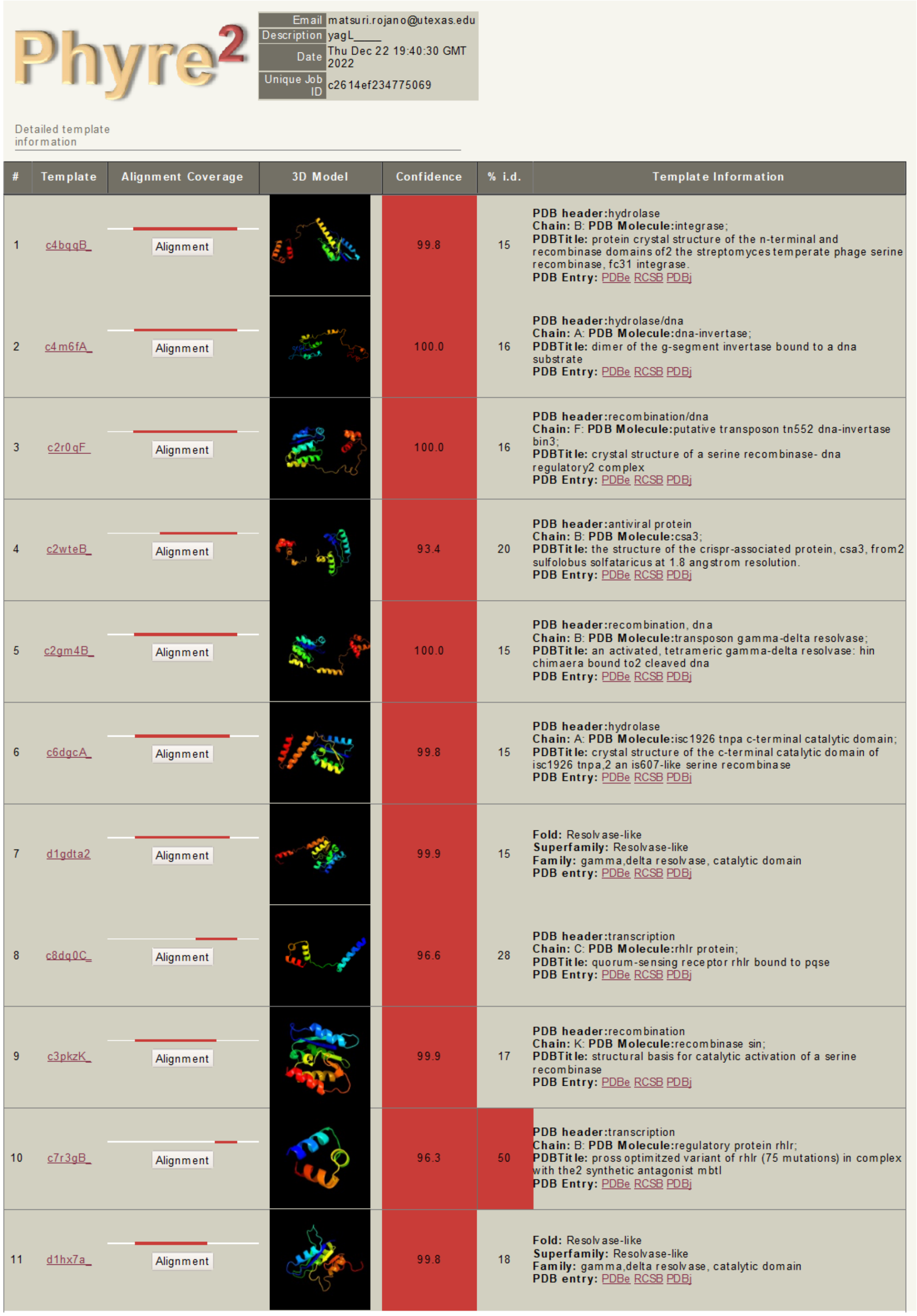

Phyre2

| | |
|---|---|
| Email | |
| Description | yagL____ |
| Date | Thu Dec 22 19:40:30 GMT 2022 |
| Unique Job ID | c2614ef234775069 |

Detailed template information

| # | Template | Alignment Coverage | 3D Model | Confidence | % i.d. | Template Information |
|---|---|---|---|---|---|---|
| 1 | c4bqqB_ | Alignment | | 99.8 | 15 | **PDB header:**hydrolase **Chain:** B: **PDB Molecule:**integrase; **PDBTitle:** protein crystal structure of the n-terminal and recombinase domains of2 the streptomyces temperate phage serine recombinase, fc31 integrase. **PDB Entry:** PDBe RCSB PDBj |
| 2 | c4m6fA_ | Alignment | | 100.0 | 16 | **PDB header:**hydrolase/dna **Chain:** A: **PDB Molecule:**dna-invertase; **PDBTitle:** dimer of the g-segment invertase bound to a dna substrate **PDB Entry:** PDBe RCSB PDBj |
| 3 | c2r0qF_ | Alignment | | 100.0 | 16 | **PDB header:**recombination/dna **Chain:** F: **PDB Molecule:**putative transposon tn552 dna-invertase bin3; **PDBTitle:** crystal structure of a serine recombinase- dna regulatory2 complex **PDB Entry:** PDBe RCSB PDBj |
| 4 | c2wteB_ | Alignment | | 93.4 | 20 | **PDB header:**antiviral protein **Chain:** B: **PDB Molecule:**csa3; **PDBTitle:** the structure of the crispr-associated protein, csa3, from2 sulfolobus solfataricus at 1.8 angstrom resolution. **PDB Entry:** PDBe RCSB PDBj |
| 5 | c2gm4B_ | Alignment | | 100.0 | 15 | **PDB header:**recombination, dna **Chain:** B: **PDB Molecule:**transposon gamma-delta resolvase; **PDBTitle:** an activated, tetrameric gamma-delta resolvase: hin chimaera bound to2 cleaved dna **PDB Entry:** PDBe RCSB PDBj |
| 6 | c6dgcA_ | Alignment | | 99.8 | 15 | **PDB header:**hydrolase **Chain:** A: **PDB Molecule:**isc1926 tnpa c-terminal catalytic domain; **PDBTitle:** crystal structure of the c-terminal catalytic domain of isc1926 tnpa,2 an is607-like serine recombinase **PDB Entry:** PDBe RCSB PDBj |
| 7 | d1gdta2 | Alignment | | 99.9 | 15 | **Fold:** Resolvase-like **Superfamily:** Resolvase-like **Family:** gamma,delta resolvase, catalytic domain **PDB entry:** PDBe RCSB PDBj |
| 8 | c8dq0C_ | Alignment | | 96.6 | 28 | **PDB header:**transcription **Chain:** C: **PDB Molecule:**rhlr protein; **PDBTitle:** quorum-sensing receptor rhlr bound to pqse **PDB Entry:** PDBe RCSB PDBj |
| 9 | c3pkzK_ | Alignment | | 99.9 | 17 | **PDB header:**recombination **Chain:** K: **PDB Molecule:**recombinase sin; **PDBTitle:** structural basis for catalytic activation of a serine recombinase **PDB Entry:** PDBe RCSB PDBj |
| 10 | c7r3gB_ | Alignment | | 96.3 | 50 | **PDB header:**transcription **Chain:** B: **PDB Molecule:**regulatory protein rhlr; **PDBTitle:** pross optimitzed variant of rhlr (75 mutations) in complex with the2 synthetic antagonist mbtl **PDB Entry:** PDBe RCSB PDBj |
| 11 | d1hx7a_ | Alignment | | 99.8 | 18 | **Fold:** Resolvase-like **Superfamily:** Resolvase-like **Family:** gamma,delta resolvase, catalytic domain **PDB entry:** PDBe RCSB PDBj |

*Top 20 template structures identified in Phyre2 for YagL modeling (pg. 1)*

| | | | | | | |
|---|---|---|---|---|---|---|
| 12 | d2gm4a2 | Alignment | 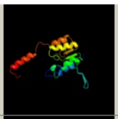 | 99.9 | 14 | **Fold:** Resolvase-like<br>**Superfamily:** Resolvase-like<br>**Family:** gamma,delta resolvase, catalytic domain<br>**PDB entry:** PDBe RCSB PDBj |
| 13 | d2rsla | Alignment | 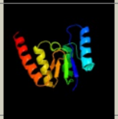 | 99.9 | 16 | **Fold:** Resolvase-like<br>**Superfamily:** Resolvase-like<br>**Family:** gamma,delta resolvase, catalytic domain<br>**PDB entry:** PDBe RCSB PDBj |
| 14 | c3lhfC | Alignment | 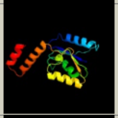 | 99.8 | 14 | **PDB header:**recombination<br>**Chain:** C: **PDB Molecule:**serine recombinase;<br>**PDBTitle:** the crystal structure of a serine recombinase from sulfolobus2 solfataricus to 2.3a<br>**PDB Entry:** PDBe RCSB PDBj |
| 15 | c2xigA | Alignment | 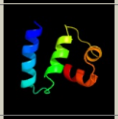 | 93.3 | 31 | **PDB header:**transcription<br>**Chain:** A: **PDB Molecule:**ferric uptake regulation protein;<br>**PDBTitle:** the structure of the helicobacter pylori ferric uptake2 regulator fur reveals three functional metal binding sites<br>**PDB Entry:** PDBe RCSB PDBj |
| 16 | c4lfuA | Alignment | 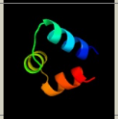 | 96.0 | 46 | **PDB header:**dna binding protein<br>**Chain:** A: **PDB Molecule:**regulatory protein sdia;<br>**PDBTitle:** crystal structure of escherichia coli sdia in the space group c2<br>**PDB Entry:** PDBe RCSB PDBj |
| 17 | c2lvsA | Alignment | 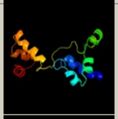 | 95.6 | 18 | **PDB header:**dna binding protein<br>**Chain:** A: **PDB Molecule:**putative uncharacterized protein;<br>**PDBTitle:** nmr solution structure of a crispr repeat binding protein<br>**PDB Entry:** PDBe RCSB PDBj |
| 18 | c3bvpB | Alignment | 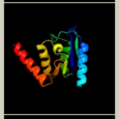 | 99.8 | 13 | **PDB header:**recombination<br>**Chain:** B: **PDB Molecule:**tp901-1 integrase;<br>**PDBTitle:** crystal structure of the n-terminal catalytic domain of tp901-12 integrase<br>**PDB Entry:** PDBe RCSB PDBj |
| 19 | c3qp5C | Alignment | 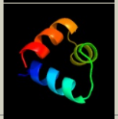 | 95.7 | 42 | **PDB header:**transcription<br>**Chain:** C: **PDB Molecule:**cvir transcriptional regulator;<br>**PDBTitle:** crystal structure of cvir bound to antagonist chlorolactone (cl)<br>**PDB Entry:** PDBe RCSB PDBj |
| 20 | c7r3eA | Alignment | 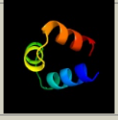 | 94.8 | 42 | **PDB header:**transcription<br>**Chain:** A: **PDB Molecule:**2-aminobenzoylacetyl-coa thioesterase,regulatory protein<br>**PDBTitle:** fusion construct of pqse and rhlr in complex with the synthetic2 antagonist mbtl<br>**PDB Entry:** PDBe RCSB PDBj |
| 21 | c4if4A | Alignment | not modelled | 95.0 | 29 | **PDB header:**transcription<br>**Chain:** A: **PDB Molecule:**response regulator protein vrar;<br>**PDBTitle:** crystal structure of the magnesium and beryllofluoride-activated vrar2 from staphylococcus aureus<br>**PDB Entry:** PDBe RCSB PDBj |
| 22 | c3ploX | Alignment | not modelled | 99.9 | 13 | **PDB header:**recombination<br>**Chain:** X: **PDB Molecule:**dna-invertase;<br>**PDBTitle:** crystal structure of the fis-independent mutant of gin<br>**PDB Entry:** PDBe RCSB PDBj |
| 23 | c2mhcA | Alignment | not modelled | 99.7 | 14 | **PDB header:**recombination<br>**Chain:** A: **PDB Molecule:**tnpx;<br>**PDBTitle:** nmr structure of the catalytic domain of the large serine resolvase2 tnpx<br>**PDB Entry:** PDBe RCSB PDBj |
| 24 | c7vimA | Alignment | not modelled | 95.7 | 38 | **PDB header:**dna binding protein<br>**Chain:** A: **PDB Molecule:**protein esrb;<br>**PDBTitle:** the c-terminal dna binding domain of esrb from edwardsiella piscicida<br>**PDB Entry:** PDBe RCSB PDBj |
| 25 | c3klnC | Alignment | not modelled | 94.0 | 41 | **PDB header:**transcription<br>**Chain:** C: **PDB Molecule:**transcriptional regulator, luxr family;<br>**PDBTitle:** vibrio cholerae vpst<br>**PDB Entry:** PDBe RCSB PDBj |
| 26 | c3sztB | Alignment | not modelled | 96.6 | 17 | **PDB header:**transcription<br>**Chain:** B: **PDB Molecule:**quorum-sensing control repressor;<br>**PDBTitle:** quorum sensing control repressor, qscr, bound to n-3-oxo-dodecanoyl-l-2 homoserine lactone<br>**PDB Entry:** PDBe RCSB PDBj |
| 27 | c6dgbA | Alignment | not modelled | 99.7 | 12 | **PDB header:**hydrolase<br>**Chain:** A: **PDB Molecule:**is607 family transposase is1535;<br>**PDBTitle:** crystal structure of the c-terminal catalytic domain of is1535 tnpa,2 an is607-like serine recombinase |

*Top 20 template structures identified in Phyre2 for YagL modeling (pg. 2)*

**Appendix 2. Additional bioinformatic analysis data for YagL modeling.**

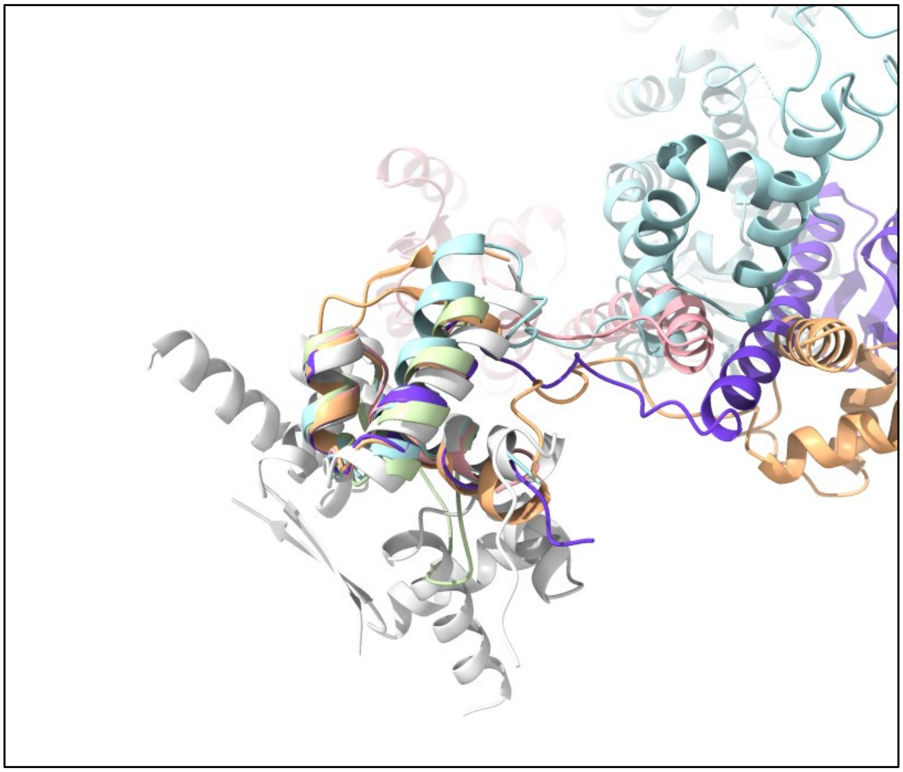

*Structural alignment of ligand protein structures to yagL HTH. Sequence-based alignment and pruned RMSD were conducted using the mmaker command in ChimeraX. Representations shown for the YagL predicted structure (purple), 1GDT (light pink), 2R0Q (light blue), 7BHY (grey), 6PAX (light orange), and 1HCR (light green).*

| Structure PDB Code | Chain | Alignment Score | Pruned RMSD (number of atom pairs; RMSD in Å) | RMSD (number of atom pairs, RMSD in Å) |
|---|---|---|---|---|
| YagL (self-alignment) | A | 237.5 | 47; 0.000 | 47; 0.000 |
| 1GDT | A | 77.2 | 33; 0.920 | 44; 2.958 |
| 2R0Q | C | 73.1 | 24; 0.893 | 43; 5.021 |
| 7BHY | C | 61.8 | 19; 0.670 | 42; 4.119 |
| 6PAX | A | 63.7 | 26; 0.651 | 47; 5.998 |
| 1HCR | A | 81.1 | 29; 1.209 | 46; 6.463 |
| 1GYT (PepA) | A | 24.3 | 6; 1.633 | 45; 18.432 |

*ChimeraX mmaker output for sequence-based alignment and superposition of ligand structure proteins with the YagL HTH domain. YagL self-alignment is shown to demonstrate maximum similarity and PepA is included to show superposition of a protein that does not contain an HTH domain.*

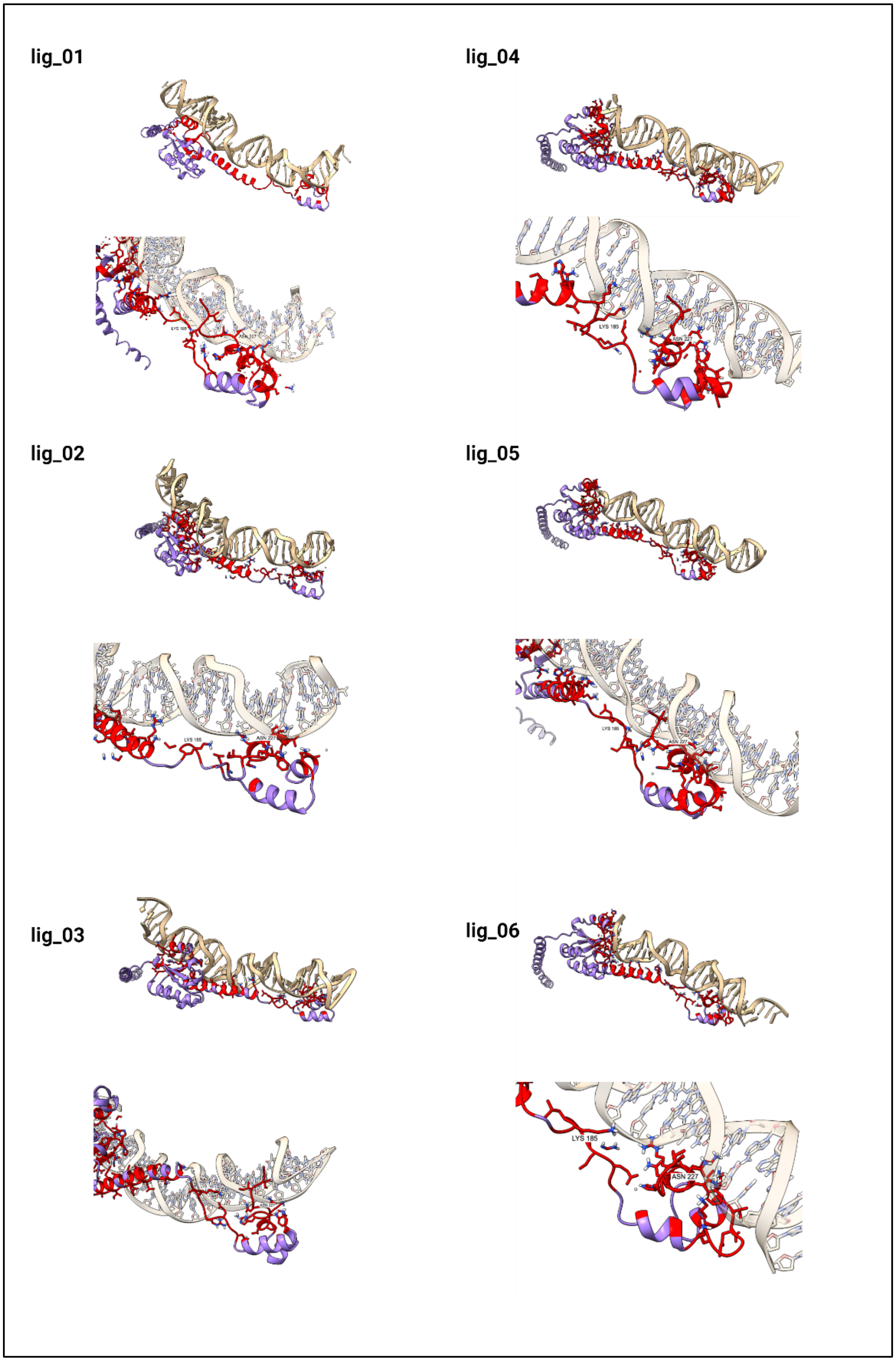


*HADDOCK docked poses for DNA ligands selected in this study.*

**Appendix 3. Gel and membrane images from *in vitro* assays.**

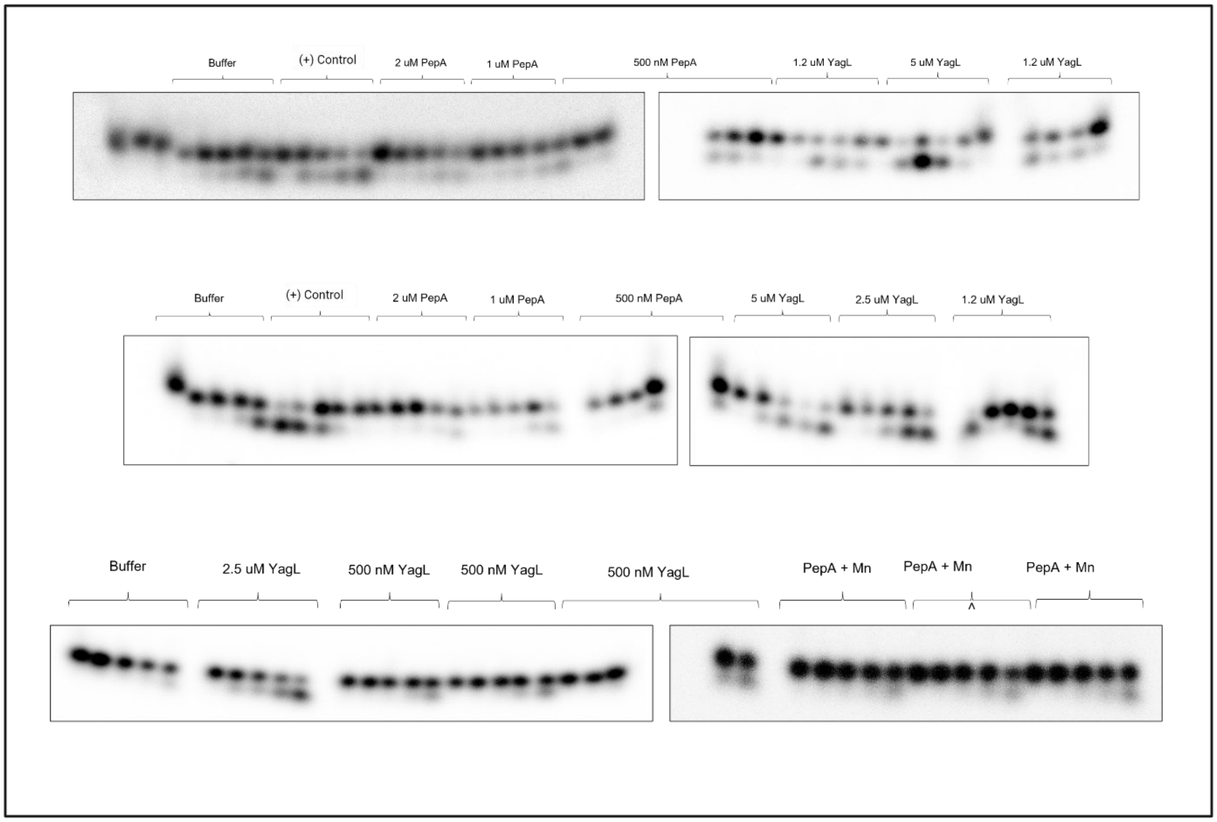


*Gel observations from the two-step catalytic activity assay (1/2).*

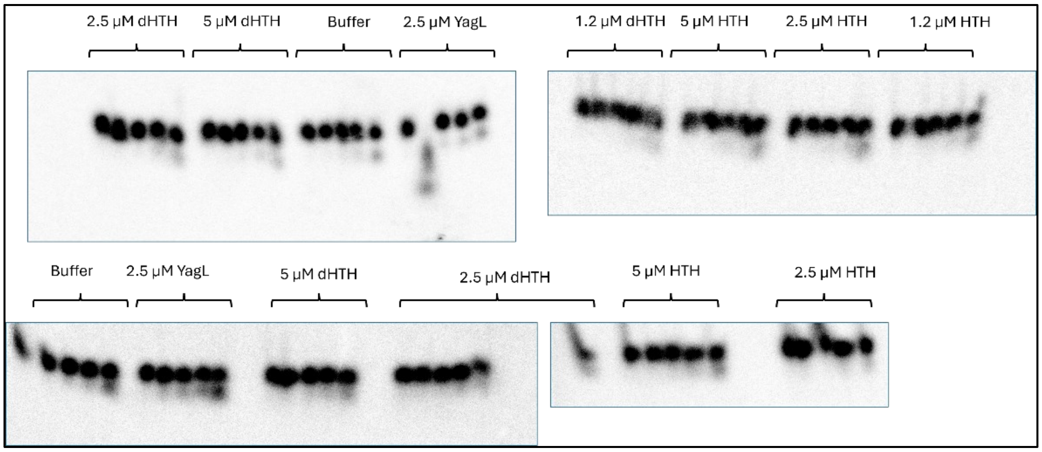


*Gel observations from the two-step catalytic activity assay (2/2).*

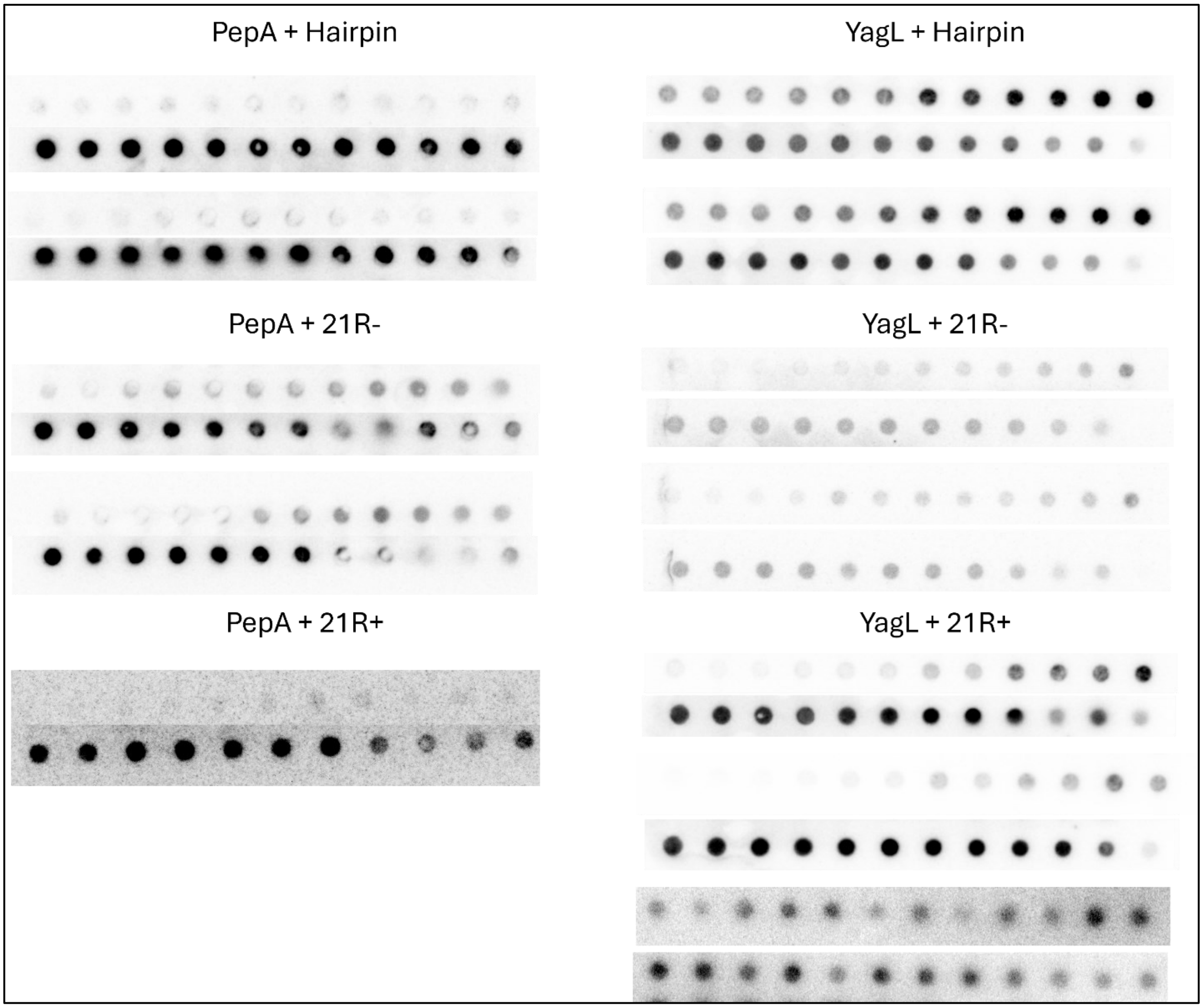


*Images from filter binding experiments (1/2).*

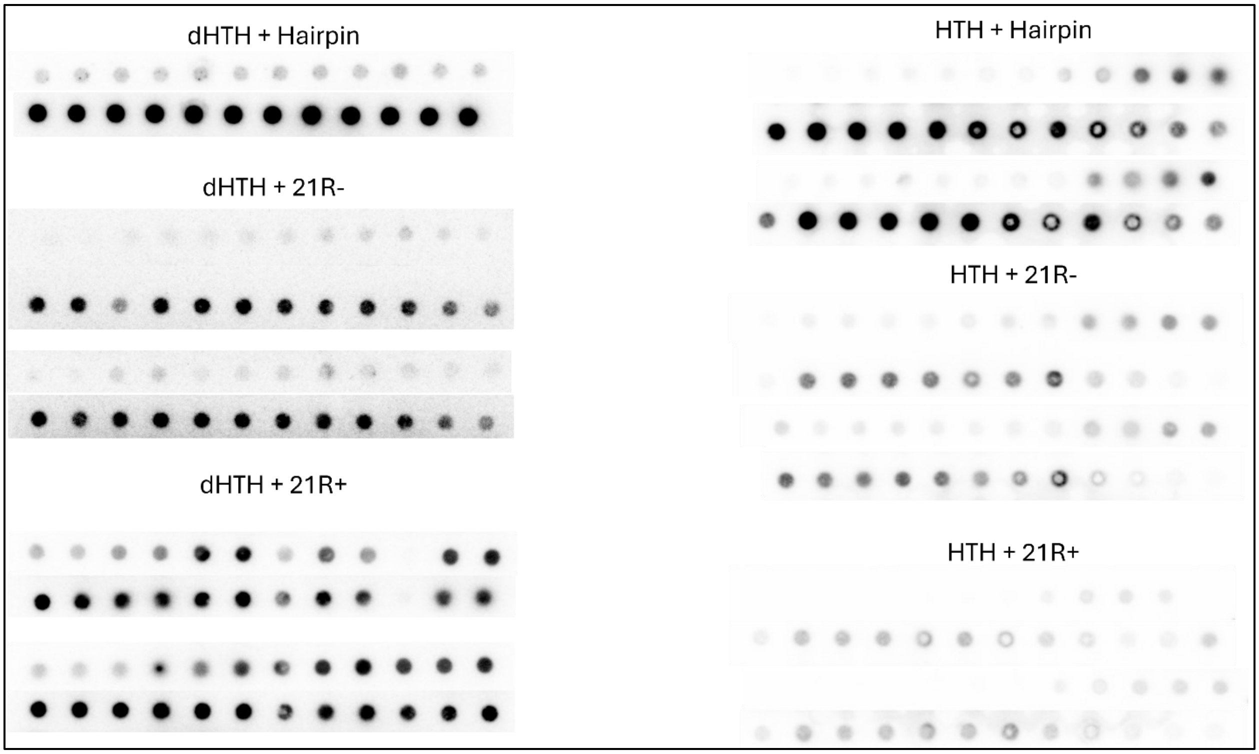


*Images from filter binding experiments (2/2).*

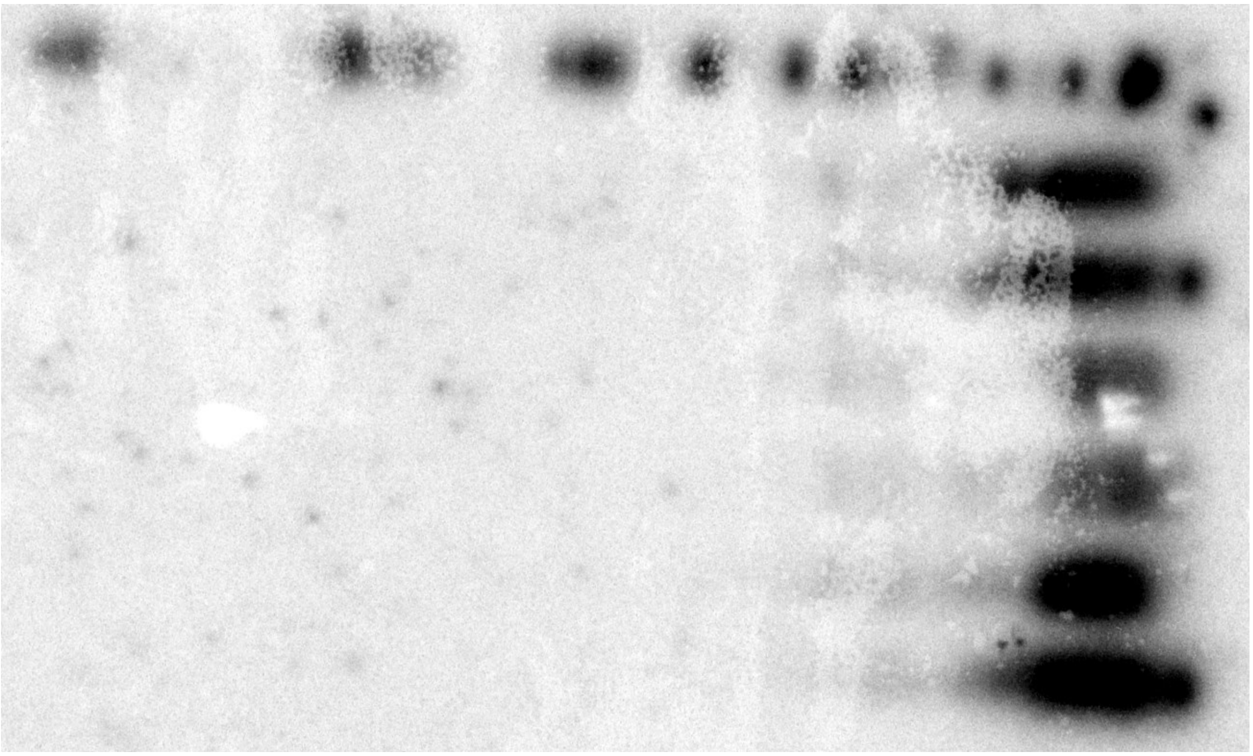

*Images from Northern Blotting experiments (1/2).*

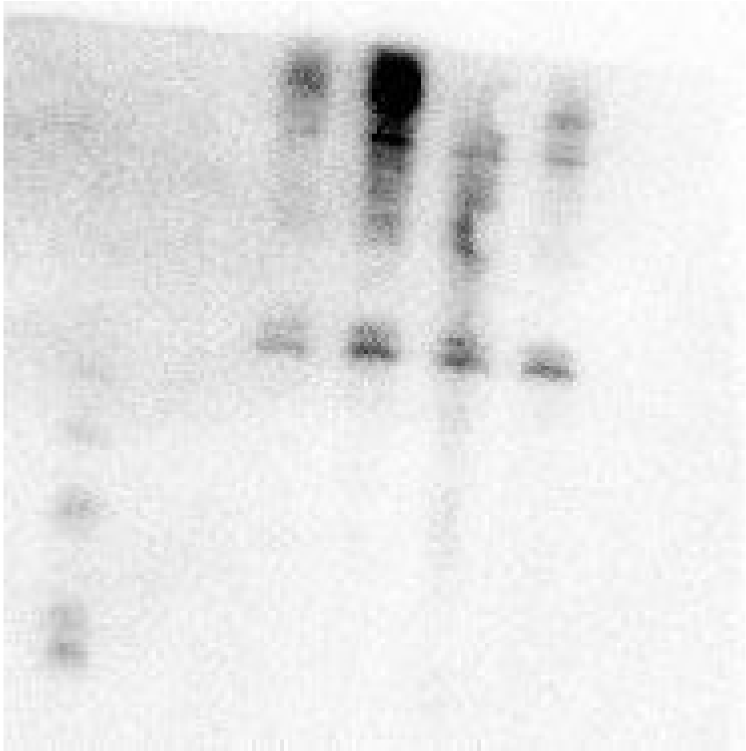

*Images from Northern Blotting experiments (2/2).*

**Supplementary References.**
